# Universal Cell Embeddings: A Foundation Model for Cell Biology

**DOI:** 10.1101/2023.11.28.568918

**Authors:** Yanay Rosen, Yusuf Roohani, Ayush Agrawal, Leon Samotorčan, Tabula Sapiens Consortium, Stephen R. Quake, Jure Leskovec

## Abstract

Developing a universal representation space for cells which encompasses the tremendous molecular diversity of cell types within the human body and more generally, across species, would be transformative for cell biology. Recent work using single-cell transcriptomic approaches to create molecular definitions of cell types in the form of cell atlases has provided the necessary data for such an endeavor. Here, we present the Universal Cell Embedding (UCE) foundation model. UCE was trained on a corpus of cell atlas data from human and other species in a completely self-supervised way without any data annotations. UCE’s modeling approach is to create a unified biological latent space that can represent cells across diverse tissues and species. This universal cell embedding captures important biological variation despite the presence of experimental noise across diverse datasets. An important aspect of UCE’s universality is that new cells can be mapped to this embedding space with no additional data labeling, model training or fine-tuning. We applied UCE to create the Integrated Mega-scale Atlas, embedding 36 million cells, with more than 1,000 uniquely named cell types, from hundreds of experiments, dozens of tissues and eight species. We uncovered new insights about the organization of cell types and tissues within this universal cell embedding space, and leveraged it to infer function of newly discovered cell types. UCE’s embedding space exhibits emergent behavior, uncovering biology that it was never explicitly trained for, such as identifying developmental lineages and embedding data from novel species not included in the training set. Overall, by enabling a universal representation for every cell state and type, UCE provides a valuable tool for analysis, annotation and hypothesis generation over single cell data.

## Introduction

Cells are the fundamental unit of life and biologists have long conceptualized cells as members of different universal landscapes [1–4]. A notable example of this is the Waddington landscape, which presents a theoretical framework for the developmental lineages of cells as they transition from pluripotent stages such as stem cells to more terminally differentiated end points [5]. Broadly, the field of cell biology has sought to map the range of phenotypes that cells might exhibit, their interrelationships, and the shifts between these states during development and disease [6–10].

The substantial growth in the size of single-cell RNA sequencing (scRNA-seq) datasets presents a fresh opportunity to revisit these questions. Detailed transcriptomic snapshots of cells are now widely available from a range of timepoints, tissues, donors, and species [11–13]. These rich, high-dimensional states are typically distilled into low-dimensional vectors or embeddings to facilitate computational analysis [14, 15]. However, existing computational approaches struggle to jointly analyze these diverse datasets. The unified representations they produce are often unable to extend to new datasets due to species-specific constraints or the presence of dataset-specific artifacts (or batch effects) which can obscure the underlying biological signal [16, 17].

Some computational methods for scRNA-seq data have managed to overcome these limitations, but at the cost of requiring model tuning for each new dataset, thus rendering the representations non-universal [15,18,19]. As a result, whenever a new experiment is performed and new data is collected, it requires dedicated, resource-intensive data labeling and model training to perform even the most standard analyses, such as clustering or annotation. This process is time consuming and inefficient, and results in sub-optimal analyses based on small, limited and private datasets.

Recent advances in the field of artificial intelligence have enabled general-purpose foundation models (such as ChatGPT [20, 21], PaLM [22], and SAM [23]) that can learn universal representations that are then applied to diverse downstream tasks and analyses. These foundation models are not specifically trained for these downstream tasks, thus presenting clear instances of emergent capabilities [24]. This foundation model strategy has also found valuable applications in biological contexts such as learning representations of protein and DNA sequences [25, 26]. Although recent work has applied foundation model architectures to single cell genomic data, the unique characteristics of these data require a specialized modeling approach to fully realize their potential [27, 28]. Directly modeling gene expression as text in the form of a sequence of genes is both inefficient from a learning perspective and often relies on inaccurate biological assumptions.

Here, we present Universal Cell Embedding (UCE), a foundation model for single-cell gene expression that is designed to address questions in cell and molecular biology. UCE is uniquely able to generate representations of new single-cell gene expression datasets with no model fine-tuning or retraining while still remaining robust to dataset and batch-specific artifacts. Moreover, it does so while requiring no cell type annotation and no input dataset preprocessing, such as gene selection. UCE can be applied to any set of protein-coding genes from any species, even if they are not homologs of genes seen during training. UCE learns a universal, intrinsically meaningful representation of cell biology that enables insights extending beyond experimentally observed data.

The representations learned by UCE display an emergent organization of cell types and states that is consistent with known biology. These cell embeddings can be used to accurately predict cell types/states without additional model retraining, showing improved performance in dataset integration against existing atlas-scale integration methods. UCE also enables the mapping of new data into a universal embedding space, already populated with annotated reference states. This strategy addresses issues such as noisy measurements that limit data alignment between different experiments and reduces the reliance on small sets of marker genes to translate insights between studies [29]. UCE can foster novel cross-dataset discoveries and overcome the limitations currently faced when working with small, isolated datasets. For example, a cell type classifier trained to predict specific immune cell types can be seamlessly applied to a completely new dataset.

## Results

### A biologically-informed foundation model for single cell gene expression

Integrating single-cell RNA sequencing (scRNA-seq) datasets is challenging for two primary reasons: scRNA-seq data does not always contain the same genes, or features, and those features are plagued by dataset-specific experimental artifacts or batch effects, which means models have to be built separately for each dataset. UCE overcomes these challenges by abstracting cells as ‘bags of RNA’ [30]. UCE (Fig. 1a) converts the RNA gene expression of a single cell into an expression-weighted sample of its corresponding genes. Next, UCE represents the sample’s genes by their protein products, using a protein language model. This allows UCE to meaningfully represent any protein-coding gene, from any species based on sequence alone, regardless of whether the species had appeared in the training data. After integrating additional metadata on gene chromosomal locations, the representation is processed by a large transformer model [31]. UCE can then map any cell—from any tissue or species—into a shared universal space without further training.

**Figure 1.**
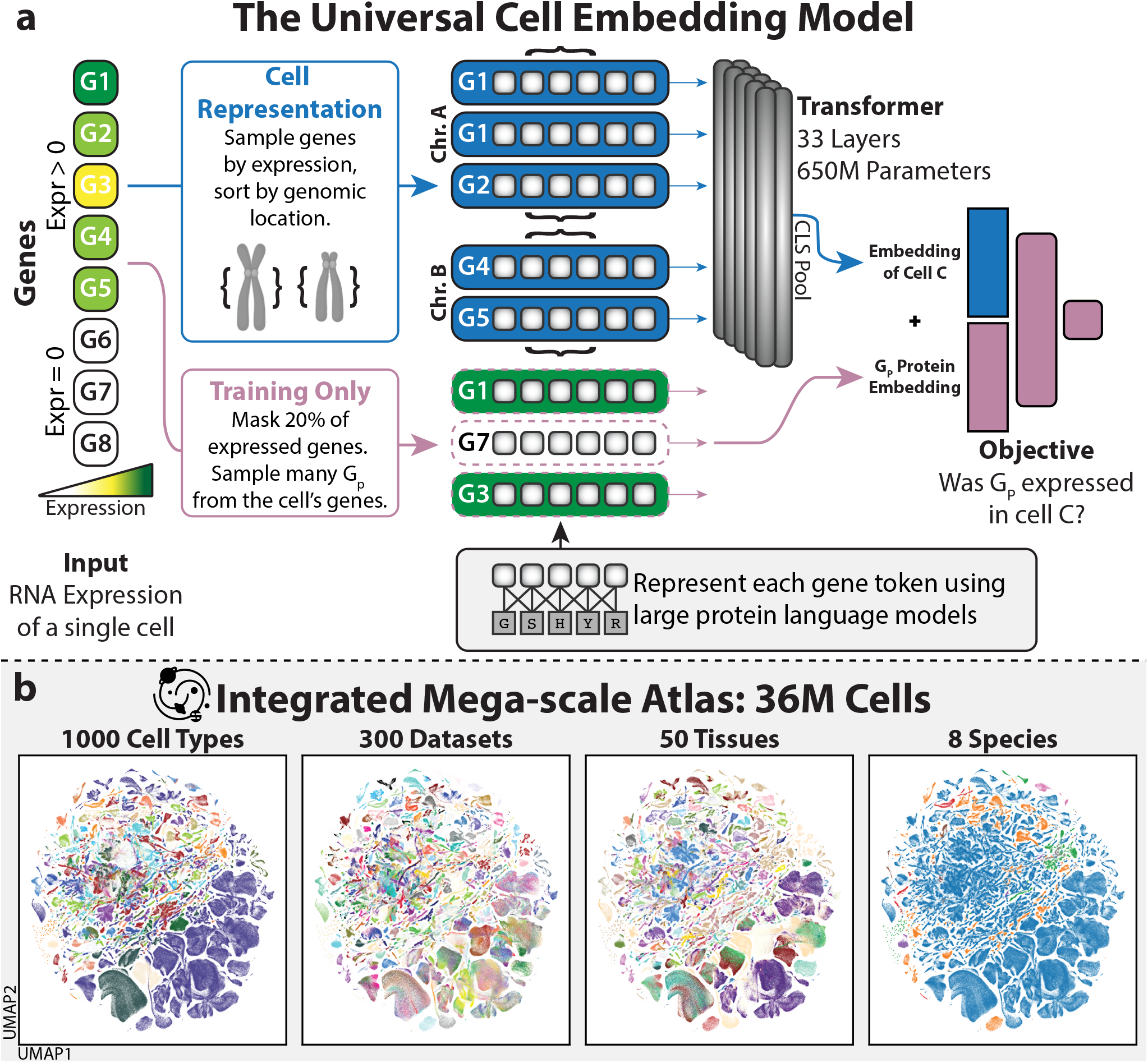
The Universal Cell Embedding Model is a large foundation model for single cell biology. **(a)** Overview of the Universal Cell Embedding (UCE) model. UCE has a unique, biologically motivated representation of cells (blue) and training scheme (purple). Given the gene expression for a single cell, UCE samples with replacement genes that were expressed, weighted by their level of expression. Each gene is represented using a ‘token’ corresponding to its protein product. Gene tokens are represented numerically by using ESM2 protein embeddings, a 15 billion parameter protein language model that takes amino acid sequences as an input. The gene tokens are sorted by genomic location and grouped by chromosome. Chromosome groups are delineated by specific chromosome start tokens and end tokens, joined, and then passed into a transformer neural network. The embedding of the cell is determined by taking the final layer output of a special CLS token that is appended before all the other tokens. To train the UCE model, a portion of genes that were expressed are masked. The model next combines the protein embeddings corresponding to each of these genes with the embedding of the cell, and passes this joint representation through a neural network that predicts if a given gene was expressed in the cell or not. This objective function is then used to update the weights of the model. **(b)** UMAP visualizations of the universal cell embedding space. We apply UCE to embed 36 million cells, with more than 1,000 uniquely named cell types, from hundreds of datasets, dozens of tissues and eight species (Supplementary Figure 19), creating an Integrated Mega-scale Atlas (IMA) spanning the universe of cell biology.

UCE takes as input (1) scRNA-seq count data and (2) the corresponding protein embeddings, generated by a protein language model, ESM2 [32], for the genes in the data set. The ESM2 protein language model takes amino acid sequences as an input and produces a numerical representation called a protein embedding. Given the expression count data for a cell, UCE takes a weighted and normalized sample, with replacement, of the cell’s genes. This sample can only contain genes which had nonzero expression and can contain multiple copies of each gene. These genes are then tokenized by converting them to the protein embedding representation of the protein that they code for [33]. Genes belonging to the same chromosome are grouped together by placing them between special tokens and then sorted by genomic location. A special token representing the entire cell, the ‘CLS’ token, is attached to the beginning of the cell representation [34]. This combined representation is passed into a transformer neural network. The embedding of a cell is taken as the embedding of the CLS token at the final layer of the transformer (Fig. 1a).

UCE is trained in a fully self-supervised manner, without using any cell type or dataset-based annotations. While incorporating cell type labels into the model design or training could seem advantageous, most datasets lack such annotations. Ultimately, it is preferable for the organization of cells within UCE to emerge organically, independent of human labeling.

UCE is a 33 layer model consisting of over 650 million parameters. UCE was trained across more than 300 datasets that are largely collected from the CellXGene corpus [35] consisting of over 36 million cells, for 40 days across 24 A100 80GB GPUs (Methods, Extended Data Table 2, SI Table 2). We also performed extensive ablation experiments to validate UCE architecture choices, understand training behavior, and quantify the benefits of ESM2 protein embeddings (SI Tables 3, 4, 8, 9, 10, Supplementary figures 17, 18). The model’s weights and implementation are freely available and the model is hosted as an openly available resource for the community.

### UCE creates an Integrated Mega-scale Atlas (IMA) of 36 million cells

We apply UCE to generate an Integrated Megascale Atlas (IMA) of 36 million cells sampled from diverse biological conditions, demonstrating the emergent organization of UCE cell representations (Fig. 1b). We find that cells within the UCE space naturally cluster under biological conditions such as cell type, while mixing among experimental conditions such as batch (Fig. 1 b). Since UCE is trained using only unlabeled data, this organization represents an emergent behavior of the model. We inspect how tissue residency can influence the state of cell types. Cells in the IMA have been prelabeled by their cell type. As these labels were never used for training the UCE model, we use them to validate the quality of the learned representation. Although macrophages found in different tissues are characterized by diverse transcriptional identities [36], they align closely in the UCE space (Extended Data Table 1). For example, human macrophages are found in 73 different tissues and among these tissues, 72% (53) of tissue-specific macrophage centroids were embedded closest to a macrophage centroid from another tissue. Similar cross-tissue homogeneity can also be identified in other prolific cell types, like endothelial cells or neurons. This shows that UCE, without explicit training or labels, identifies that macrophages have a unique cellular identity that is shared between tissues. More broadly, it is an example of a UCE’s emergent organization that is consistent with known biology, even though not explicitly trained for.

### UCE embeds new datasets without additional model training

We evaluated the quality of UCE representations by directly mapping new datasets which were not part of the training set into the embedding, without any additional training or refinement of the UCE model. This is called a ‘zero shot’ capability, since the model was never trained on any samples from the new dataset (Fig. 2a). While a variety of deep learning models have been proposed for this cell embedding task, we choose to compare the performance of UCE to other self-supervised transformer-based methods. This is because they do not rely on cell type annotation and are trained on large datasets. In particular, we compare against Geneformer [27] and scGPT [28]. Geneformer represents cells as lists of genes sorted by their expression, while scGPT represents cells as lists of genes with binned expression values.

**Figure 2.**
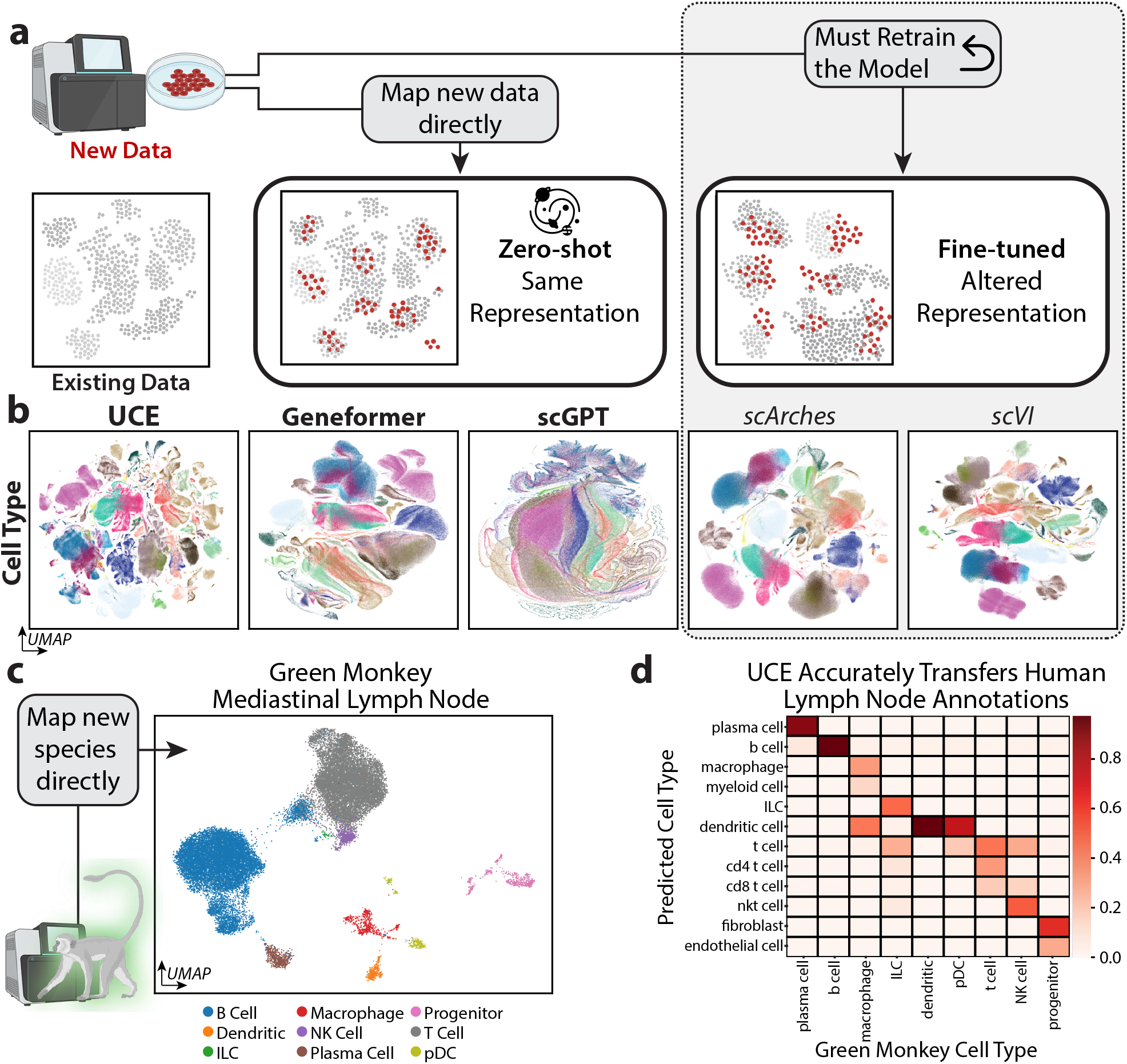
Zero-shot cell embedding capabilities of UCE. **(a)** Comparison of zero-shot and fine-tuned single-cell embedding models. A zero-shot embedding model maps new data directly to the representation space, with no additional model training. In contrast, fine-tuned models must first be retrained on a given dataset, and only then can be applied on that dataset, fundamentally altering the model’s representation space. **(b)** UMAP embeddings of UCE and other methods for Tabula Sapiens v1 and v2, colored by cell type. UCE zero-shot embeddings outperform other methods when scored using metrics from the single cell integration benchmark (Supplementary Table 1) [16]. **(c)** UMAP of cells from a new species, green monkey colored by cell type. UCE is able to generate high-quality zero-shot embeddings of novel species that were never seen during training. The UCE embedding for green monkey mediastinal lymph node [40] recaptures cell type clusters. Notably, a population of cells annotated as B cells (blue) clusters nearby to T cells. This breakaway cluster of putative T cells expresses canonical T cell markers including *Cd3, Fyb1, Il7r, Hcst* and *Gzmk* (Supplementary Fig. 3). **(d)** Green monkey lymph node cells can be accurately annotated using the IMA. A logistic classifier is first trained to predict cell types based on UCE embeddings of human lymph node cells. The classifier is then directly applied on green monkey cells to predict the cell types. Predicted cell types have high agreement with the original cell type annotations, demonstrating that UCE can be used to transfer cell type annotations to novel species.

We assess the performance of these methods on a completely new and yet unreleased dataset (as of the training of the model), Tabula Sapiens v2, which contains diverse human data from 581,430 cells, 27 tissues, 167 batches, and 162 unique cell types. Tabula Sapiens is collected in a highly standardized way and represents one of the largest ‘gold-standard’ expert annotated cell atlases available. We use established metrics for embedding quality that measure the conservation of cell type information and the correction of batch effects (Methods). We found that UCE substantially outperforms the next-best Geneformer method by 13.9% in the overall score, 16.2% in the biological conservation score, and 10.1% on batch correction score (SI Table 1). UCE embeddings distinctly separate cell types more effectively than other methods (Supplementary fig. 1, 2). To further evaluate the value of UCE’s zero-shot embeddings, we compare them to fine-tuned methods that rely on cell type labels and dataset-specific training. Remarkably, UCE performs slightly better than these established approaches, including scVI [15] and scArches [18] (SI Table 1).

Tabula Sapiens v2 includes cells measured using both droplet or plate-based sequencing methods. Thus correcting technology-based batch effects is a difficult task. UCE zero-shot embeddings of Tabula Sapiens v2 Ovary tissue, which contains 45, 757 cells profiled using 10x-primev3 and 3, 610 cells profiled using Smart-seq3, successfully correct batch effects at the same level as fine-tuned methods, while more accurately representing cell types. When scored using the single cell integration benchmark (SCIB), UCE’s batch correction scores are close to that of scVI and scArches, while its bio conservation scores are higher (SI Table 5). Furthermore, we demonstrate how UCE can simply transfer cell-type labels. We train a simple logistic classifier on the UCE embeddings of the Immune Cell Atlas [37], and then apply the classifier to cell embeddings from Tabula Sapiens v2. We observe that the classifier accurately classifies Tabula Sapiens v2 cells as memory and naive B cells (Supplementary Fig. 2b), which is confirmed by marker gene analysis (Supplementary Fig. 2c).

Beyond Tabula Sapiens v2, we evaluate UCE on held-out (not seen during training) complex datasets with hundreds of cell types [12, 38] and on an additional large-scale benchmark that measures how well embedding spaces capture cell-type relationships across samples in million-cell datasets with hundreds of cell types and batches [11, 12, 39], finding that UCE outperforms Geneformer and scGPT and matches the performance of fine-tuned methods such as scVI (Supplementary Note 6; Supplementary Tables 10 and 12).

### UCE embeds diverse cell types from organisms that were not part of the training data

UCE can also align datasets from previously unseen species without additional training. This capability arises because UCE is genome-agnostic: each gene is translated into its corresponding protein sequence and embedded within a universal protein space. As a result, the representation is species-independent and does not rely on identifying homologous gene pairs. Because UCE can analyze cell atlas data from species outside its training set, its performance on this task provides a stringent test of how well its emergent representations capture underlying biological principles.

UCE’s training data is composed of datasets from eight species: human, mouse, mouse lemur, zebrafish, pig, rhesus macaque, crab-eating macaque and western clawed frog (Supplementary Figure 19). We apply UCE to embed datasets from three novel species that were not included in the training set. For each species, we generate a zero-shot embedding and then determine the nearest cell type centroid from the IMA for each of the dataset’s existing annotated cell types.

Within a dataset of green monkey lymph node and lung cells [40], for 13 of the 17 cell type centroids, the closest centroid from another species corresponds to the same cell type in the green monkey. This match extends to all 17 centroids when considering the three nearest centroids (Extended Data Table 1, Fig. 2c, 2d). Moreover, a population of lymph node cells that were originally labelled as B cells, form a distinct cluster in UCE space (Supplementary Fig. 3b). Differential expression analysis revealed that this cluster predominantly expresses T cell markers, *Cd3d, Fyb1, Il7r, Hcst* and *Gzmk* (Supplementary Fig. 3a, 3c) [41, 42].

In the case of naked mole rat spleen and circulating immune cells [43], for 17 of 24 cell types, the nearest cross-species centroid matches the naked mole rat cell type (Extended Data Table 1, Supplementary Fig. 4b). In the case of chicken, we embed two different chicken data sets, the chick retina [44] and the developing chick heart [45] (Supplementary figs. 5a, 5b). Different eye-specific neurons within the chick retina map to mouse lemur neurons, such as chick oligodendrocytes, which are closest to mouse lemur oligodendrocytes (Extended Data Table 1). In chicken heart, 12 of 15 cell type centroids are matched within the nearest two cross species centroids (Extended Data Table 1). No bird species were included when training UCE.

We compare UCE’s performance to a state-of-the-art supervised method for cross-species label transfer, SATURN [33, 46] and SAMap [19]. For each of the four datasets, we integrate them with a paired human dataset from the same tissue from Tabula Sapiens v1, except for chicken retina, which was paired with retina data from Orozco et al [47]. In comparison to UCE, which is zero-shot and unsupervised, SATURN is a supervised method: it is aware of each species’ cell type annotations individually. Both SATURN and SAMap are finetuned methods, trained from scratch on the human and novel species dataset and designed explicitly to map cell types together. Still, UCE outperforms both methods for cell type label transfer between the human and the new species for three out of four datasets, (Supplementary Note 5, Supplementary Table 14). UCE produces coherent zero-shot embeddings even for highly divergent species such as Drosophila [48] (Supplementary Fig. 20). At the same time, fine-grained cell-type transfer across such large evolutionary distances remains challenging, reflecting both reduced cross-species alignment performance and inherent ambiguity in defining reliable “ground-truth” correspondences between distantly related cell-type ontologies (Extended Data Table 4).

### UCE learns a meaningful organization of cell types in previously unseen data

Moving beyond metrics focused on individual cell type clusters, we also examined the structure of the universal embedding space as a whole, through the relative positioning of different cells within it. A meaningful arrangement of cell types emerges upon embedding of all cells from the Tabula Sapiens v2 dataset from lung tissue (Fig. 3a). Not only do distinct cell types like T cells, monocytes and endothelial cells cluster together, but higher-level categories, such as immune cells and epithelial cells, are also clearly distinguished. When compared to the cell hierarchy derived using the Cell Ontology [49], UCE identified clusters showed greater similarity (measured using the adjusted Rand index) than other zero-shot embedding methods (Supplementary fig. 14).

**Figure 3.**
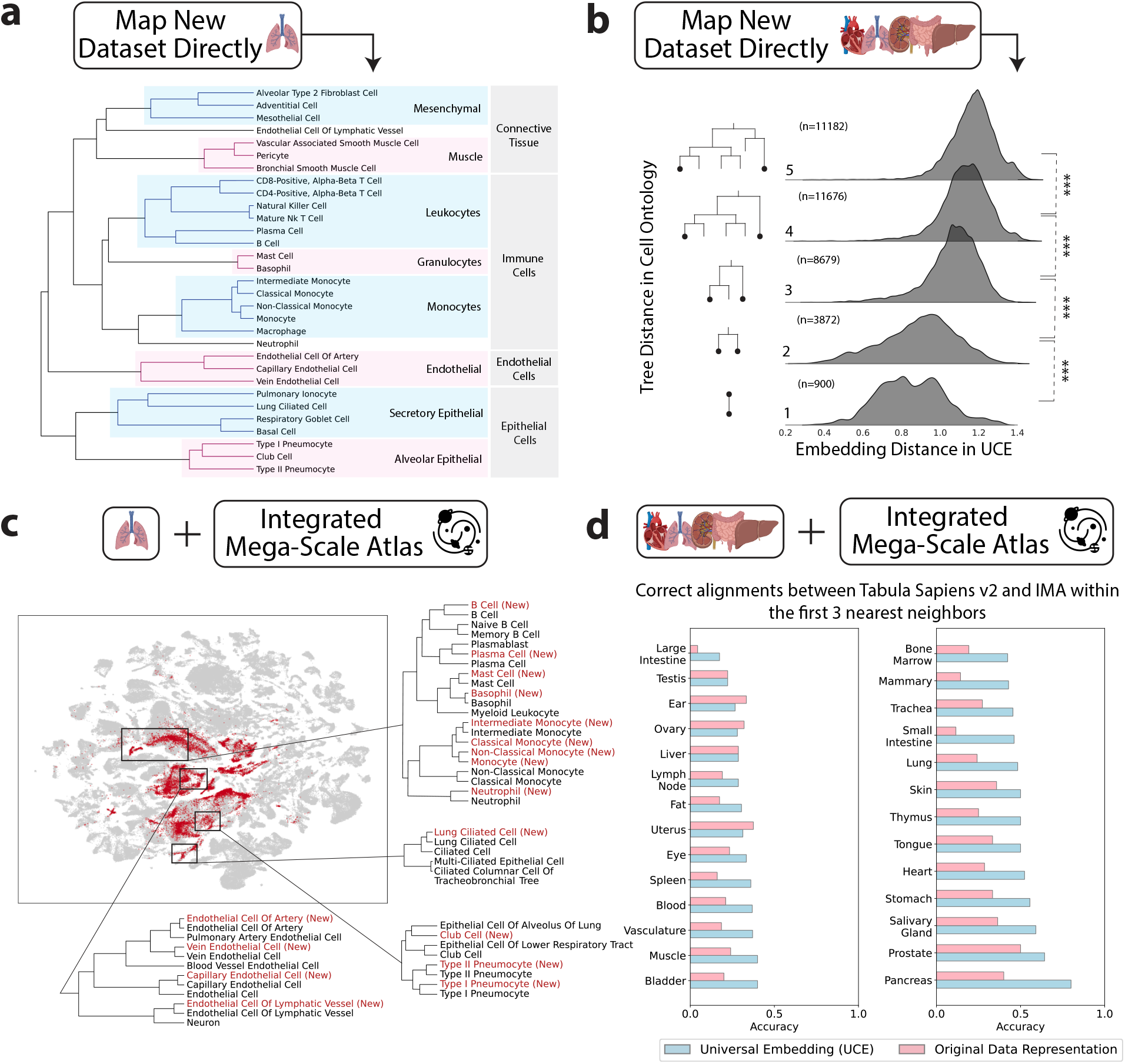
UCE learns meaningful organization of cell types. **(a)** The UCE space generated for new, previously unseen data shows a meaningful arrangement of cell types. Lung data was used from new donors from the Tabula Sapiens Consortium. Dendrogram of hierarchical clustering of all annotated cell types in the UCE embedding space. Closely connected cell types in the dendrogram show meaningful relationships both at finer and coarser scale resolutions. **(b)** Evaluation of the organization of cell types in the embedding space when compared to Cell Ontology. The *x*-axis depicts the density of Euclidean distances between all pairs of cells across all tissues for these new donors from the Tabula Sapiens Consortium. The *y*-axis shows the corresponding tree distance between cell types as found in the Cell Ontology. Stars denote statistical significance, which was established using a one-sided *t*-test, p values in increasing order of distance: 10^−106^, 10^−201^, 10^−25^, 10^−131^ **(c)** Mapping data from new donors to the Integrated Mega-scale Atlas (IMA) across multiple lung datasets. Red labels correspond to data from new donors, grey are from IMA datasets. All cell type labels from multiple datasets are displayed as-is, with no modifications or reformatting of text. Accurate alignment between the new dataset and IMA is observed at finer resolution. Four different subtypes of endothelial cells are shown to correctly map to their corresponding counterparts in the complete mega-scale atlas. In the case of lung ciliated cells, they map more closely to their matching counterpart as compared to all other ciliated cell subtypes also present in the IMA. **(d)** Quantification of cell type alignment between new dataset and IMA. Accuracy in 3-nearest centroid matches between new dataset and IMA cell types at the finest level of original annotation. Results are measured across all 27 tissues in Tabula Sapiens v2 for both the UCE space and the original gene expression space. Tissues are ordered by accuracy in the UCE space.

To further assess the organization of all cells within the embedding, we compared distances between pairs of cell types across all tissues in the embedding space to their distances in the Cell Ontology tree (Fig. 3b). We hypothesized that cells that are known to be similar based on the cell ontology would likely also be closer together in the embedding space, and that the degree of closeness would be correlated with ontological similarity. The results validate this relationship: at each additional unit of separation between cell types in the cell ontology tree, there is a significant increase in the embedding distance in UCE between those cell types (Fig. 3b). We observed this trend up to 5 hops in the ontology tree, after which it plateaued (Supplementary fig. 6), likely due to the curse of dimensionality and variations in ontological refinement across branches (SI Note 3).

The precision of the organization of cell types in the Tabula Sapiens v2 lung dataset was evaluated by comparing it with other lung datasets in the IMA (Fig. 3c, Supplementary Fig. 8). Four different endothelial cell subtypes are observed to map correctly to their corresponding counterparts in the IMA (Fig. 3c). Similarly, lung ciliated cells correctly map to their counterpart in the larger corpus despite the presence of four different subtypes of ciliated cells. Further analysis of the alignment of cell-type centroids between Tabula Sapiens v2 and the IMA in all tissues showed an exact alignment for 41% of cell types (Methods). This alignment, based on the three nearest neighbor cell type centroids, is 65% more accurate compared to that measured in the original gene expression space (Fig. 3d). When focusing on the nearest centroid, the alignment accuracy improves by 92%. Many inexact matches are caused by differences in the labeling resolution of different datasets in the IMA (SI Tables 6, 7, Extended Data Table 8). These results demonstrate that UCE can effectively learn a universal representation of cell biology that not only enables discrimination between individual cell types but also captures their relative similarities across scales with the potential to reveal deeper insights into development and function.

### A workflow for decoding the function of newly discovered cell types

UCE’s zero-shot embedding capabilities unlock novel computational analyses of scRNA-seq data and aid in hypothesis generation. UCE differs from other methods in that the same cell type can also be compared against all previously assayed cells across tissues, disease states and species. UCE is not biased by existing annotations, opening the door to discovery of a novel function (Fig. 4a). With existing fine-tuning-based methods, every searched dataset would need to be integrated, requiring repeated model retraining. Thus, UCE enables a new workflow for scRNA-seq data analysis that performs an unbiased search across the universe of cell biology.

**Figure 4.**
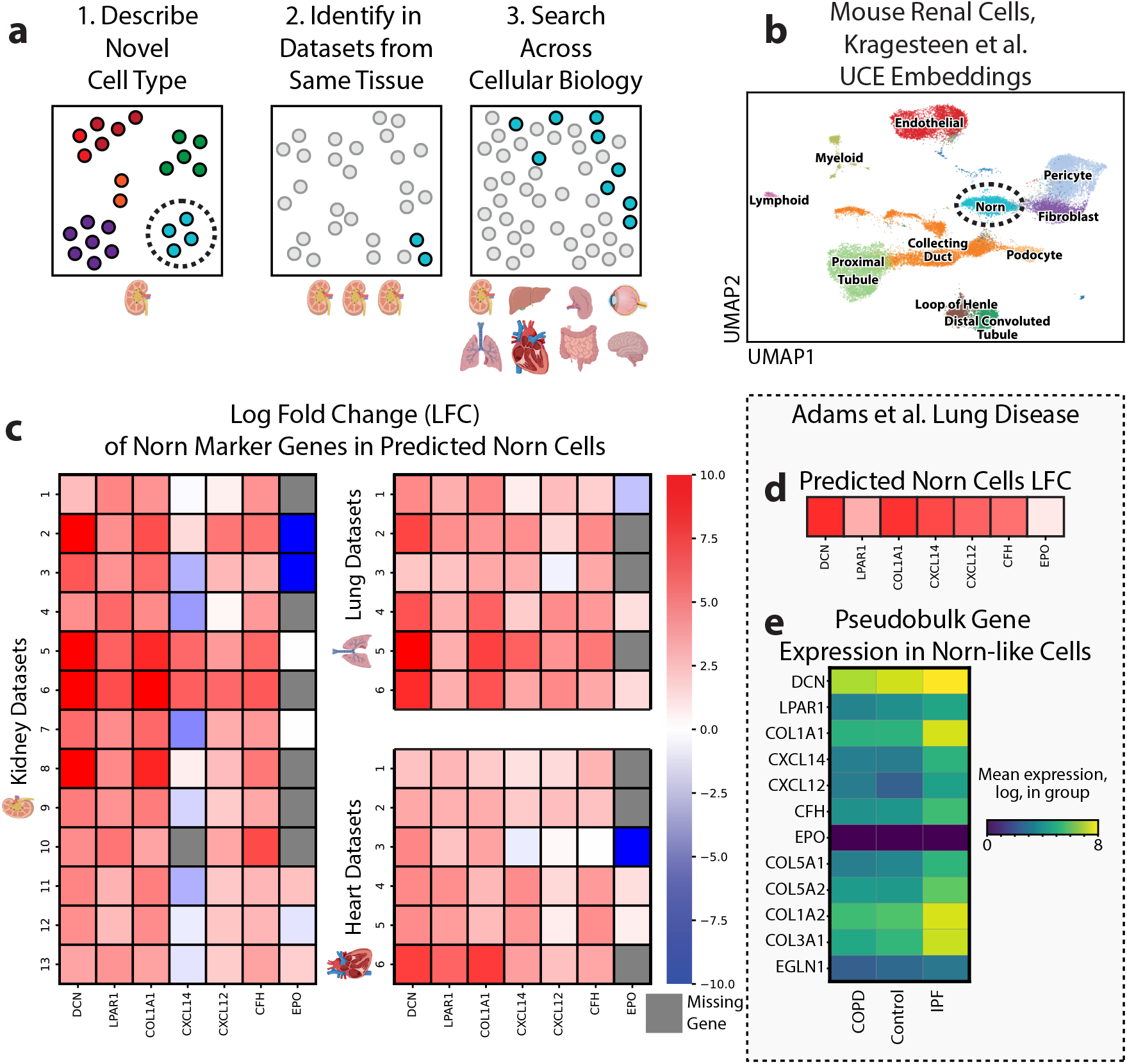
Norn Cell Case Study: UCE unlocks new analyses of single cell datasets. **(a)** Overview of a novel single cell analysis workflow that UCE facilitates. Analysis begins with (1) the identification of a novel cell type (circled) within the embedding space, using methods such as clustering and confirmation using marker gene analysis. (2) Next, the novel cell type can be easily identified in other datasets profiled from the same tissue (for example, kidney). A simple classifier, such as a logistic classifier, is trained to predict cell types from universal cell embeddings, and is then applied to embeddings from other datasets of the same tissue (kidney), to confirm the cell type’s existence and improve its characterization. (3) Finally, the same simple classifier can be applied to the embeddings of cells from any other tissue, to find cell types with similar biological functions or patterns of gene expression. **(b)** Identification of novel Norn cells in mouse kidney. UMAP visualization of zero-shot embedding of mouse renal cells from Kragesteen et al. [50]. Norn cells form a distinct cluster within the embedding space (circled), and a logistic cell type classifier trained on the dataset recaptures Norn cell labels 98.3% of the time. **(c)** Identification of Norn cells and Norn-like cells across tissues. A logistic classifier is trained to predict Norn cells from universal cell embeddings, and is then applied to other kidney datasets (left) and datasets from lung and heart (right). The log fold change of known Norn marker genes between cells predicted to be Norn cells and the remaining cells within each dataset is visualized. Cells which are predicted to be Norn-like preferentially express Norn markers in kidney, as well as in lung and heart. Notably, *Cxcl14* has a mixed pattern of expression among some datasets. **(d)** Cells predicted to be Norn-like cells within a lung disease dataset [52] express known Norn markers, as demonstrated by log fold change (LFC). **(e)** Pseudobulk differential gene expression (Supplementary Table 13) in predicted Norn-like cells, grouped by disease status. Norn markers have similar expression levels across disease status, except for *Cxcl14*, which has significantly higher expression in IPF versus COPD or control, and *Cxcl12*, which is expressed more highly in IPF versus control. Norn-like cells in IPF patients preferentially express genes encoding collagen compared to COPD patients (*Col1a1, Col5a1, Col1a2, Col3a1*). *Egln1* is expressed at a lower rate in IPF norn-like cells compared to COPD norn-like cells.

We present an example of this analysis by using the recently identified kidney Norn cell as a case study. The kidney Norn cell is the long-sought erythropoietin (*Epo*) producing cell in the kidney, and is characterized as fibroblast-like. We perform a zero-shot embedding of mouse renal cells from [50], which produces a cluster of cells corresponding to Norn cells (Fig. 4b).

Using a simple logistic classifier trained on the embedding of mouse renal cells, we predict the existence of Norn cell clusters in many kidney datasets. Since this classifier takes universal cell embeddings as input, we can directly apply it to all 36 million cells in the IMA, in a manner unbiased by cell-type annotations ascribed by previous studies. To evaluate the model’s predictions that these are Norn, or Norn-like cells, we investigate the expression of canonical Norn cell marker genes. Cells classified as Norn cells in the top 13 kidney datasets by Norn abundancy demonstrate preferential expression of the Norn markers *Dcn, Lpar1, Col1a1, Cxcl12*, and *Cfh* (Extended Data Table 3). Notably, *Epo* transcripts, which are often missing from data sets and are lowly expressed, are not typically differentially expressed in these cells. *Epo* cannot readily be used as a marker gene for Norn cells because of a variety of factors, including low overall expression, even in hypoxic Norn cells, the bursty nature of *Epo* transcription, and the fast degradation of *Epo* messenger RNA upon reoxygenation [50]. *Cxcl14*, another marker of Norn cells, displays mixed expression patterns in these predicted Norn cells (Fig. 4c). The same pattern of marker gene expression is also found in cells from other tissues, including lung and heart datasets (Fig. 4c). Furthermore, these predicted cells also share a common set of genes that are lowly expressed in mouse renal Norn cells (SI, Fig. 9). The tissues with the highest number of predicted Norn cells were gonad, heart and lung. Finally, we also identify differences in gene expression specific to each tissue (Supplementary fig. 10). While *Epo* expression has been previously observed in the heart and lung tissue, the mechanisms and cell types associated with this expression, and their relation to kidney Norn cells have not been previously determined [51].

### UCE helps interrogate alternate lung disease outcomes

Lastly, we apply UCE and our Norn cell classifier to investigate Norn-like cells in lung diseases. We generate an embedding of lung cells sampled from patients with idiopathic pulmonary fibrosis (IPF), chronic obstructive pulmonary disease (COPD), or patients from a control group [52]. We identify Norn-like lung cells that preferentially express Norn markers in all three groups (Fig. 4d). For these Norn-like lung cells, we identify differences across disease groups (Fig. 4e). COPD and IPF are both associated with elevated bloodstream *Epo*, but COPD has levels higher than IPF. Furthermore, in patients with IPF, secondary erythrocytosis is absent or reduced compared to patients with COPD [53–55]. Given the identification of Norn-like cells in the lung, and Norn cell’s production of *Epo*, it is possible that this difference in disease prognosis could be related to disease associated differences in Norn-like cells.

IPF predicted norn-like cells do not have significantly different expression of norn marker genes compared to COPD patients, except for *Cxcl14*, which they express at a significantly higher rate. However, they do preferentially express collagen genes (*Col1a1, Col5a1, Col1a2, Col3a1*), and have significantly lower expression of the gene *Egln1*, which encodes the oxygen sensing enzyme PHD2 (Fig. 4e, Supplementary Table 13) [50].

Taken together, these results indicate that cells with similar transcriptional states to Norn cells can be found in other tissues in the body, and such cells may play a previously undescribed role in disease. UCE greatly facilitates an analysis of this scale and diversity because it is a universal model that is agnostic to tissue, species or disease state. In order to ground this analysis, results are confirmed using expression of canonical marker genes (Supplementary Fig 9) .

## Discussion

UCE is a single-cell foundation model that is built from the ground up to represent cell biology across the wide array of single-cell datasets. We envision UCE as an embedding approach that enables researchers to map any new data, including entire atlases, into an accurate, meaningful and universal space. The embedding space that emerges from UCE is highly structured and diverse and aligns cell types across tissues and species. Additionally, these cell types organize themselves in a pattern that reflects existing biological knowledge.

By building UCE, we enable novel analyses of scRNA-seq data. UCE’s zero-shot embeddings are an important new capability because they enable an intrinsically meaningful representation that can extend insights beyond the data that have already been observed and annotated experimentally. Our results demonstrate that UCE can achieve such a generalizable representation across different datasets while maintaining accuracy comparable to methods that require retraining for each specific dataset. To ensure that UCE’s embeddings, or any downstream models trained on them, remain universally shareable, we have designed UCE to be used as-is, without any model fine-tuning. This allows UCE representations to serve as a universal way of connecting any single-cell gene expression dataset.

While UCE represents a significant advance toward universal cell embeddings, several caveats temper its promise and highlight directions for improvement. Current benchmarks primarily evaluate UCE’s ability to recover expert-annotated cell types, but these tasks are inherently limited by the granularity and subjectivity of available cell labels. As such, they may not fully capture the model’s capacity to represent subtler biological processes, such as cellular responses to perturbations or cross-modality integration. Moreover, like other large-scale biological foundation models, UCE operates largely as a “black box,” making it difficult to interpret how or why certain embeddings arise. Addressing this opacity will require the development of new interpretability tools that connect learned representations to mechanistic biological understanding. UCE’s training data, while extensive, is biased toward mammalian species—particularly human and mouse—and to specific tissues such as brain, raising concerns about its generalizability to underrepresented species and contexts. Methodological improvements in dataset construction could help alleviate these biases while reducing computational burden. Finally, although UCE improves alignment across datasets by capturing shared biological signal, it does not replace traditional batch effect correction methods, which remain useful for handling known experimental confounders. Similarly, its reliance on gene sampling based on expression means that some fine-grained quantitative variation is lost, potentially limiting the resolution of certain cell state distinctions.

In 2002, Nobel laureate Sydney Brenner identified many of the core motivations for the creation of cell atlases and virtual cells. Virtual cells should be the goal of biological foundation modeling, because cells are the “real units of function and structure in an organism” [56]. Brenner also identified the need for such models to be computationally efficient, predictive, and able to generate new cell types. We believe that UCE represents a significant advancement in the progress towards a virtual cell [57]. Through learning a universal representation of every cell state and type, we expect that UCE will be a valuable tool for analysis, annotation and hypothesis generation as the scale and diversity of single-cell datasets continues to grow.

## Methods

### Overview of UCE

UCE (Universal Cell Embedding) is a machine learning model for mapping single-cell gene expression profiles into a universal embedding space, denoted as 𝒰. In this space, each cell *c*_*i*_ is represented as a *d*_*emb*_-dimensional vector, where *d*_*emb*_ = 1280. For all analyses we use the 1280 dimensional UCE representation and not a 2 dimensional embedding such as that generated from tSNE or UMAP. Users of UCE should rely on the full 1280 dimensional representations to judge distances between cells, rather than data visualization using these methods, especially when they expect to see strong biological differences.

The model takes as input a dataset 𝒟 with *N* cells 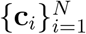. Cells in 𝒟 can be drawn from one or more distinct scRNA-seq experiments. Each cell *c*_*i*_ in 𝒟 is described by a gene expression vector 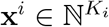, where *K*_*i*_ is the number of genes measured in *c*_*i*_ and can differ across 𝒟. The gene expression vectors 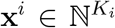 are not subset to those with high variance. UCE defines a function 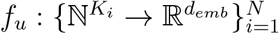 that maps each gene expression vector **x**^*i*^ to its cell embedding vector **h**^*i*^.

### Model input: Gene representation

The expression of gene *g* in cell *c*_*i*_ is denoted by 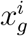, where *g* represents any protein-coding gene. The corresponding token embedding *p*_*g*_ is a pretrained embedding for the protein(s) encoded by the gene *g*. These embeddings are derived from a pretrained protein language model that takes an amino acid sequence as input and returns a *d*_*p*_-dimensional embedding vector as output. To create *p*_*g*_, we take the average of all proteins coded by gene *g*. In the context of UCE, we can formulate this as a dictionary that maps each gene *g* to a *d*_*p*_-dimensional protein embedding vector. Specifically, we employ the ESM2 model, which yields embeddings of size *d*_*p*_ = 5120 [32, 33].

Protein language models are chosen for gene representation because they can generate universal representations of any protein sequence. Therefore, for new species, all that is required is to have the amino acid sequence of that species’ protein coding genes. These genes do not need to have solved structures and orthology does not need to be calculated between them and the existing training data. Model training ablations demonstrate that among a class of different protein language models, UCE models trained with ESM2-15B performed the best (Supplementary Fig. 18). Model ablation experiments further demonstrate that the use of protein embeddings to tokenize genes has significant benefits for the more poorly represented species in the training corpus, as a small ablation model trained with protein embeddings outperformed a model with randomly initialized embeddings on all species except for human (Supplementary Fig. 18). Additionally, the use of protein embeddings enables a unique capability: embedding new species not found in the training data (Fig. 2.)

### Model input: Cell representation

For each cell *c*_*i*_ in the input dataset 𝒟, we identify two distinct sets of protein-coding genes: the expressed genes 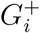 and the non-expressed genes 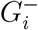. These sets are defined as follows:

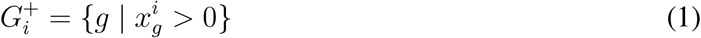

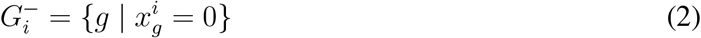

For producing the cell embedding, a multi-set of 1024 non-unique genes 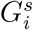 are sampled from the expressed genes 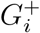, with replacement. The probability of sampling a gene 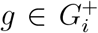 is weighted by the log normalized expression of that gene, which can be formulated as:

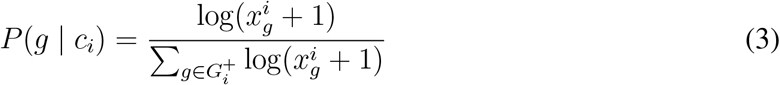

where 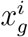 is the expression count of gene *g* in cell *c*_*i*_, and the sum in the denominator is over all genes in 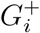.

Note that for a given cell, different random seeds can yield different sampled gene subsets and thus slightly different representations (Supplementary Table 11); however, using a fixed seed selects the same genes each time and produces an identical embedding for repeated runs on the same cell.

Once the multi-set 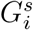 is compiled for each cell *c*_*i*_, we arrange the genes within each chromosome according to their genomic positions. Different chromosomes are specified using special chromosome start and end tokens. Start tokens are unique to each chromosome and species. Every chromosome group is combined into a single sequence, with chromosome order randomly determined. A cell-level *CLS* token is appended to the start of the sequence. It is designed to capture the cell-level embedding upon training the model. The final sequence of genes ordered by genomic location and separated by chromosome is referred to as the cell sentence *S*_*i*_ for cell *c*_*i*_.

Ordering the genes using this information does appear to have a positive effect on model performance (Supplementary Note 4, Supplementary Table 3). Transformer models do not require any ordering of their inputs, and this metadata was only added to the model in order to enable potential applications, such as examining chromosomal aberrations or synteny between species. Future models of this type may choose to exclude this metadata, or include it using other methods besides positional encodings.

### Transformer Architecture

Each cell sentence *S*_*i*_ is fed into a transformer that consists of *n*_*lay*_ layers. Each layer contains a multi-head self-attention mechanism with *n*_*head*_ attention heads and a feedforward network operating over a hidden space of dimensionality *d*_*hid*_. We also initialize sinusoidally-varying positional embeddings. Gene token embeddings are compressed using a single layer MLP to *d*_*emb*_-dimensional vectors before passing through the transformer.

While the transformer architecture is highly performative, it requires significant computational cost to train and evaluate. One important aspect of this cost is the quadratic increase in the runtime of transformers’ attention operations proportional to the number of tokens. Such a relationship requires that the number of genes sampled by UCE, 1024, be relatively limited, especially given that individual cells might have more than 1024 uniquely expressed genes. Future model architectures could explore increasing this context length, or using models with sub-quadratic scaling such as state space models [58]

### Model output: Cell embedding

The final output from the model is the cell embedding vector 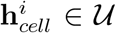 which corresponds to the *d*_*emb*_-dimensional embedding of the *CLS* token in the final layer of the model following decoding with an additional MLP.

When cells with unrealistic random gene expression patterns, such as those generated by shuffling the expression of real cells, are inputted, the resulting cell embeddings default to a heterogeneous, out of distribution output (Supplementary Fig. 11).

### Model training: Cell representation

At the time of training, we generate a set 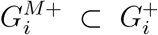 by randomly selecting a certain percentage (*r*_*mask*_) of genes from 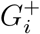, without replacement. This set is used for computing the loss during training, and is masked from the cell representation.

The probability of sampling a gene 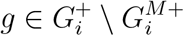 (Equation 3) is then updated to be:

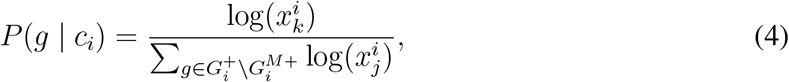

We also establish two additional gene sets to be used for loss computation: 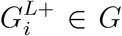 and 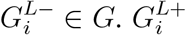 and 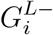 are randomly selected from the masked set of expressed genes 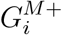 and the set of unexpressed genes 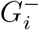 respectively. Both 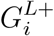 and 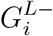 are of equal size, specifically *N*_*loss*_*/*2. In the case of 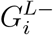, the sampling is done without replacement unless 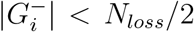. Similarly 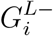, is also sampled without replacement unless 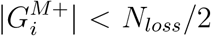. In this case, 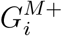 is used as-is along with additional samples drawn with replacement from the full set of expressed genes 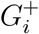.

### Model training: Loss Function

UCE is trained to reconstruct the binarized expression of genes in a cell, when those genes are masked from the model by setting their expression to 0. To calculate the loss function for a given cell *c*_*i*_, the cell embedding vector 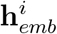 is individually concatenated with every gene *g* within both 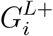 and 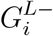. These concatenated vectors then serve as input to a feedforward multilayer perceptron (MLP), which computes the probability that gene *g* is expressed within cell *c*_*i*_.

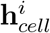 represents the embedding vector for cell *c*_*i*_ and *p*_*g*_ represents the token embedding for gene *g*. Then the concatenated vector 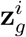 that serves as input to the MLP for cell *c*_*i*_ and gene *g* is:

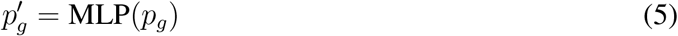

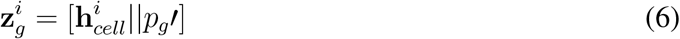

where || denotes the concatenation operation and 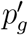 is the compressed protein embedding.

The MLP then processes this concatenated input to produce the predicted probability that gene *g* is expressed:

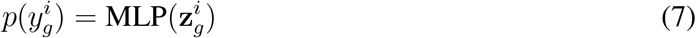

This probability is then used in the binary cross-entropy loss function. The true classification labels for each gene’s expression status in cell *c*_*i*_ are represented by the vector **y**^*i*^. UCE is trained to accurately predict the expression of genes in 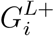 and the lack of expression in 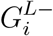. The model is trained using a binary cross-entropy loss, which is averaged across all *N*_*loss*_ genes and all *N* cells in the minibatch as follows:

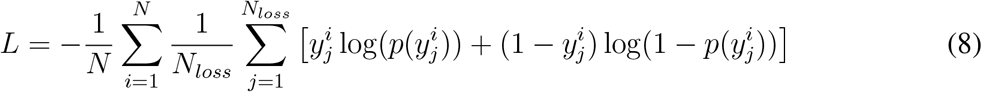

For further details on hyperparmeter choices please see Supplementary Table 2. Different hyperparameter choices for this masked loss can impact final model quality. Reducing the masking percentage from 20% to 10% or increasing it to 40% did not improve model performance (Supplementary Note 4, Supplementary Table 4). In future models, an alternate loss, for example, a generative decoder style loss, could be used, and might improve performance since no genes would be masked during training.

### Creating the IMA and dataset preprocessing

The Integrated Mega-scale Atlas (IMA) used to train UCE was created by combining scRNAseq datasets from multiple publicly available sources. The majority of IMA data (33.9 million cells and 285 datasets) is human and mouse data downloaded from CZI Cell X Gene (CxG) Census [35] version “2023-07-10” (July 10th, 2023). Duplicate cells were removed by selecting primary cells only. The remainder of the IMA is composed of 2.3 million cells from 28 datasets, from eight different species: human, mouse, zebrafish, rhesus macaque, crab-eating macaque, mouse lemur, frog, and pig.

For datasets from the CxG Census, preprocessing only involved filtering cells by minimum gene counts (200) and genes by a minimum cell counts of 10. No highly variable gene selection was applied. For datasets collected from other sources, preprocessing was not uniform (Supplementary Note 1).

For visualization of the IMA (Fig. 1b), predicting green monkey cell types (Fig. 2d), matching new species centroids (Extended Data Table 1), and prediction of Norn-like cells (Fig. 4, Supplementary Fig. 9) a representative sample of the IMA was used in place of the full 36 million cells. This representative sample was used in order to speed up computationally intensive tasks like UMAP calculation. The sample was created by randomly choosing 10,000 cells from each dataset, without replacement. For datasets with fewer than 10,000 cells, the entire dataset was included. In total, this representative sample has 2,969,114 cells. The average number of cells per dataset in the sample is 9486. For visualization and centroid calculation, cell types in the sample were coarsened by mapping them to a set of 51 coarse cell types (Extended Data Table 5).

### Model Evaluation

- **Zero-shot embedding quality and clustering** For evaluating the quality of embeddings, we used metrics from the single-cell integration benchmark [16].
- **Cell type organization** For each cell type dendrogram the Euclidean distance was used to perform hierarchical clustering across all cells.
- **Comparison to cell ontology** Here, we used the tree distance between any two cell types in Cell Ontology [49]. We used the most up to date version of Cell Ontology at the time of writing this paper (Release date: 2024*/*08*/*16). To determine the Euclidean distance distribution, we sampled 100,000 random pairs of cells from Tabula Sapiens v2.
- **Zero-shot cell type alignment to IMA** For each cell type *θ*, a centroid was identified separately for data from Tabula Sapiens v2 (TSv2) 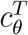 and from IMA 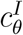. For each cell type that is present in both TSv2 and the IMA, the 3 nearest neighbor cell type centroids 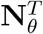 to the centroid in Tabula Sapiens 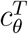 were identified. These neighbors could be either from Tabula Sapiens or from the IMA. If this set of neighbors 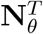 to the anchor centroid from TSv2 data 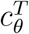 contains the centroid for the same cell type in IMA data 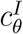, then this was counted as a correct match. This analysis was performed per tissue, both in the UCE embedding space as well as in the original expression space (after log-normalization). In case of the original data representation, the set of 5704 shared genes across all 184 human datasets in the Integrated Mega-scale Atlas (IMA) were used to represent each cell. The expression counts in each cell are normalized by total counts over all genes, so that every cell has the same total count after normalization. This is followed by a log transformation. This follows standard Scanpy data preprocessing guidelines [59].
- **Scanpy Analyses and Package Versions** Unless otherwise stated, all analyses were done using Python versions 3.7+ and scanpy version 1.93+. Default settings were always used unless otherwise specified. For differential expression analyses, the scanpy function sc.pp.highly variable genes was used with default settings with the flavor seurat v3. Scanpy was also used to generate all umaps. First, sc.pp.neighbors was run, using default settings except for setting use rep to the correct embedding value, such as the embedding from UCE. Then sc.tl.umap was used with default settings to generate the umap. The evaluation framework for UCE is released with the following versions: numpy 1.26.4, scipy 1.14.1, pandas 2.2.2, tqdm 4.66.5, torch 2.1.1, scanpy 1.10.2, accelerate 0.24.0, and for downloading model files: requests 2.25.1, urllib3 1.26.6.

### Description of other models

- **Geneformer** Geneformer [27] is a transformer based foundation model. Geneformer represents a cell as a list of genes sorted by their expression. Geneformer was trained using masked language modeling, on 30 million cells. For all analyses, the *GF-12L-30M-i2048* model was used.
- **tGPT** tGPT [60] is a transformer based foundation model. tGPT represents cells, like Geneformer, as a list of genes sorted by expression. However, tGPT is trained as an autoregressive langauge model, rather than a masked langauge model. tGPT was trained on 22.3 million cells.
- **scGPT** scGPT [28] is a transformer based foundation model. scGPT represents cells as a list of the expression values of its genes. scGPT is trained to iteratively decode those genes’ expressions when some are masked, using generative pretraining. scGPT was trained on 33 million cells. For all analyses, the *whole-human* model was used.
- **scVI** scVI [15] is a variational autoencoder. scVI is trained to reconstruct gene expression values using a zero-inflated negative binomial loss. scVI was used with default parameters for all experiments. scVI version 1.01+ was used. scVI input datasets were transformed only by taking the highly variable gene subset using sc.pp.highly variable genes with the flavor seurat v3, and then the scVI model was trained with default settings (number hidden variables 10).
- **scArches** scArches builds on top of the scVI architecture to enable transfer learning of cell types. As used in this work, scArches first trains an scVI model on a reference dataset, fine-tunes a cell-type aware scANVI model using the cell types from the reference dataset, and then further finetunes this model on a transfer dataset using architecture surgery. scArches was used with default parameters for all experiments. scArches version 0.5.1 was used, and was trained using the same scVI model for the dataset as a starting point.
- **PCA** For comparisons to PCA, default scanpy settings and best practices were used. The dataset was normalized using sc.pp.normalize total which normalized cell counts to the median count value for the dataset. Expression values were then log normalized using sc.pp.log1p, and pca was calculated with sc.pp.pca, which defaults to the top 50 principal components.

### Differential expression analysis of predicted Norn cells

A logistic classifier was trained to predict cell types from UCE embeddings on mouse kidney cells. This classifier was then applied to UCE embeddings from the representative sample of IMA datasets. Datasets were then split by tissue, and the datasets with the most predicted norn cells in each tissue were used for differential expression analysis. The top 13 kidney datasets, top 6 lung and top 6 heart datasets were chosen.

For each individual (full) dataset, RNA counts were log normalized, and then differential expression was run using default settings as implemented in Scanpy [59], comparing predicted Norn cells to all other cells in the dataset. The results of these differential expression tests were used to determine the log fold change of marker genes in predicted Norn cells (Fig. 4c, Supplementary Fig. 9).

## Data availability

The full list of datasets used to train UCE are in Extended Data Table 2. Most of these datasets are available to download from CellXGene [35]. Tabula Sapiens v2, used for model evaluation, will be made available upon publication.

Datasets analyzed in the paper are publicly available to download. The green monkey lung and lymph node dataset is available with accession code GSE156755. The naked mole rat dataset is available with accession code GSE132642. The chicken retina dataset is available with accession code GSE159107. The chicken heart dataset is available with accession code GSE149457. The mouse kidney dataset is available with accession code GSE193321. The human lung disease dataset is available with accession code GSE136831.

## Code availability

UCE was written in Python using the PyTorch library. The source code is available on Github at https://github.com/snap-stanford/uce Code for reproducing figures and analyses is available on Github at https://github.com/yhr91/uce_reproduce/tree/master.

## Acknowledgements

We thank Rok Sosič, Kexin Huang, Charlotte Bunne, Hanchen Wang, Michihiro Yasunaga, Michael Moor, Minkai Xu, Mika Jain, George Crowley, Maria Brbić, Jonah Cool, Nicholas Sofroniew, Andrew Tolopko, Ivana Jelic, Ana-Maria Istrate and Pablo Garcia-Nieto for discussions and for providing feedback on our manuscript. We acknowledge support from Robert C. Jones for help with accessing and analyzing the Tabula Sapiens v2 dataset. We acknowledge support from the Chan Zuckerberg Initiative, including help with accessing and processing CxG datasets. We gratefully acknowledge the support of DARPA under Nos. N660011924033 (MCS); NSF under Nos. OAC-1835598 (CINES), CCF-1918940 (Expeditions), Stanford Data Science Initiative, Wu Tsai Neurosciences Institute, Amazon, Genentech, GSK, Hitachi, Juniper Networks, and KDDI. Y. RH. acknowledges funding support from GlaxoSmithKline. L.S. was supported by the American Slovenia Education Foundation (ASEF). Icons created with BioRender.com.

## Author information

Y.RS., Y.RH., S.Q. and J.L. conceived the study. Y.RS, Y.RH., S.Q. and J.L. performed research, contributed new analytical tools, designed algorithmic frameworks, analyzed data and wrote the manuscript. Y.RS. and Y.RH. performed experiments and developed the software. A.A. and L.S. contributed to code and performed analyses. T.S. provided annotated data.

