## Supplementary Information for "Universal Cell Embeddings: A Foundation Model for Cell Biology"

### Supplementary Notes

1. **Supplementary Note 1**: Datasets and preprocessing

2. **Supplementary Note 2**: Hyperparameters

3. **Supplementary Note 3**: Comparison to Cell Ontology

4. **Supplementary Note 4**: Model Ablation Experiments

5. **Supplementary Note 5**: Comparison to SATURN and SAMap

6. **Supplementary Note 6**: Complex Dataset Benchmarking

7. **Supplementary Note 7**: Evaluation of Cell Type Organization in UCE space

### Supplementary Figures


1. **Supplementary Figure 1**: UCE outperforms existing methods for integrating Tabula Sapiens v2
2. **Supplementary Figure 2**: UCE embeddings recapture B cell identities in Tabula Sapiens v2
3. **Supplementary Figure 3**: UCE captures *Cd3* expression in green monkey lymph node
4. **Supplementary Figure 4**: UCE can embed novel species that it was not trained on (green monkey, naked mole rat) in a zero-shot setting
5. **Supplementary Figure 5**: UCE can embed novel species that it was not trained on (chicken) in a zero-shot setting
6. **Supplementary Figure 6**: Relationship between Embedding Distance and Ontological Similarity
7. **Supplementary Figure 7**: Cell types show organization in embedding space based on developmental lineage
8. **Supplementary Figure 8**: Mapping lung data from Tabula Sapiens v2 to multiple lung datasets from the Integrated Mega-scale Atlas (IMA)
9. **Supplementary Figure 9**: Predicted Norn cells express canonical Norn Markers
10. **Supplementary Figure 10**: Differential Expression Between Norn-like cells Across Tissues.
11. **Supplementary Figure 11**: Embedding of Fake Cells with UCE
12. **Supplementary Figure 12**: Using Cell Ontology to derive Reference Lung Cell Type Organization
13. **Supplementary Figure 13**: Dendrogram of Cell Type Organization created by different cell embedding methods
14. **Supplementary Figure 14**: Evaluation of Cell Type Organization in UCE space

15. **Supplementary Figure 15**: Hierarchical clustering of cell types in the Large Intestine in the UCE space identifies developmental relationships

16. **Supplementary Figure 16**: Hierarchical clustering of cell types in the Bone Marrow in the UCE space identifies developmental relationships

17. **Supplementary Figure 17**: Model and Dataset Size Ablations

18. **Supplementary Figure 18**: Protein Embedding Tokenization Ablations

19. **Supplementary Figure 19**: Training Data Species Distribution

20. **Supplementary Figure 20**: Zero Shot Fly Cell Atlas Embedding

### Supplementary Table

1. **Supplementary Table 1**: UCE Performance on single-cell Integration Benchmark

2. **Supplementary Table 2**: UCE Model Hyperparameters

3. **Supplementary Table 3**: Model Ablation: Chromosome Positional Encoding

4. **Supplementary Table 4**: Model Ablation: Gene Masking Rate

5. **Supplementary Table 5**: Performance on Tabula Sapiens v2 Ovary Tissue

6. **Supplementary Table 6**: Tabula Sapiens v2 Cell Type Alignments to IMA for Large Intestine Tissue

7. **Supplementary Table 7**: Tabula Sapiens v2 Cell Type Alignments to IMA for Prostate Tissue

8. **Supplementary Table 8**: Homolog Gene Performance for Primary Motor Cortex

9. **Supplementary Table 9**: Homolog Gene Performance for Embryonic Limb Atlas

10. **Supplementary Table 10**: Performance on Human Brain Cell Atlas

11. **Supplementary Table 11**: Effect of Gene Sampling on Cell Embedding Similarity

12. **Supplementary Table 12**: Performance on scGraph Benchmark

13. **Supplementary Table 13**: Norn Cells Pseudobulk Differential Expression

14. **Supplementary Table 14**:Novel Species Label Transfer Performance

### Supplementary Note 1 Datasets and preprocessing

We downloaded publicly available count matrix files with cell type annotations (see data availability). For the naked mole rat data, only cells from female samples were chosen, to align with the naked mole rat female proteome. For all analyzed datasets, expression matrices were filtered down so that only cells with cell type annotations were present.

A large amount of the training data was downloaded from the CellXGene (CxG) Census API. Remaining datasets were downloaded from different sources including GSE (Extended Data Table 2). For datasets from CxG, preprocessing only involved filtering cells by minimum gene counts (200) and genes by a minimum cells count of 10. Data downloaded from other sources was filtered to 8,000 genes using the SeuratV3 Highly Variable genes method implemented in Scanpy.

### Supplementary Note 2 Hyperparameters

The model size for UCE was chosen based on the availability of computational resources. The size and number of transformer embedding layers was chosen to be the same as the ESM1b protein language model[1]. UCE model and training parameters are listed in Supplementary Table 2. UCE uses GELU[2] for all MLP layers' activation functions. All MLP layers use LayerNorm normalization[3]. All transformer layers are implemented using the default PyTorch implementation, which uses ReLU activation and applies LayerNorm activation after attention and feedforward operations. During training, the learning rate was increased linearly for the first epoch, and then exponentially decayed for the remaining 7 epochs.

### Supplementary Note 3 Comparison to Cell Ontology

Distance in cell ontology [4] was measured as the tree distance between any two cell types. To determine the Euclidean distance distribution, we sampled 100,000 random pairs of cells from Tabula Sapiens v2. While the distance between entities in cell ontology is directly correlated with UCE embedding distance for more similar cell type pairs in the Cell Ontology tree, for larger distances we see a levelling off of this effect (Supplementary Figure 6). This could be attributed to a common challenge in high-dimensional spaces called the 'curse of dimensionality' which results in most points becoming uniformly distant from each other due to the sparsity of the data in this space [5]. In addition to this, quantifying similarity in the Cell Ontology tree becomes less precise at larger distances since some branches of the tree contain more finely resolved cell types than others.

### Supplementary Note 4 Model Ablation Experiments

We perform a comprehensive set of ablations on model design and hyperparameters. In order to perform these experiments while limiting computational expense, we use a smaller version of the UCE model, implemented using a new training framework, PyTorch Lightning, rather than HuggingFace Accelerator, and train all models on the same training setup of one machine with 8 A100 GPUs. This base version of the model has $4$ layers, with $8$ attention heads, no dropout, embedding dimension $512$ and hidden dimension $2048$ and is trained on all data for $8$ epochs. All model ablations use the same batch size, $24$ cells per GPU, and do not accumulate gradients, resulting in an effective batch size of $192$. All ablations are trained using the same learning rate scheduler, which linearly increases the learning rate to a max value of $0.0003$, before decaying it using a cosine scheduler, except in the cases of models that were larger than the base model and had the max value reduced to $0.0001$ because of issues with divergence. The base model has a total of $22.1$ million trainable parameters, with $12.6$ million parameters specifically in the transformer layers. For each ablation, only one change to the base model was made.

Overall, increasing model size and training for longer improved performance (Supplementary Figure 17). Protein language model tokenization of genes, particularly the ESM2 15B parameter model, improved performance on all non-human species (Supplementary Figure 18). Inclusion of chromosome and genomic location information, encoded via positional encoding, improved performance (Supplementary Table 3), as did the choice of a 20% gene masking rate (Supplementary Table 4).

For ablations which trained the model on a reduced set of data, we tried to ensure that the new, smaller dataset had both the correct number of cells and of datasets. To do so, we performed $10,000$ random selections of either 157 (for the 50% data) or 79 (for the 25% data) of the original $313$ datasets. We then choose the data split that came closest to the correct total number of cells. The 50% dataset has $18,119,211$ cells total out of $36,238,416$, and the 25% dataset has $9,059,574$. Both smaller datasets have $4$ out of the $8$ species.

For ablations that changed the size of the embedding dimension of the model, the hidden dimension was also changed, and it was set to $4$ times the new embedding dimension.

### Supplementary Note 5 Comparison to SATURN and SAMap

We benchmark SATURN [6], SAMap [7] and UCE for the task of transferring labels between the four new species datasets and paired human datasets for a similar tissue. SATURN and SAMap are both fine tuned methods, trained from scratch on the pair of datasets and designed explicitly to force cell types together. SATURN additionally has access to cell type information for each species, and was designed specifically to force cell types from different species together. Because of this, we expect them to outperform UCE for this task. However, we find that UCE outperforms both SATURN and SAMap on three out of four of the datasets each (Supplementary Table 14).

To benchmark label transfer performance, we paired each new species dataset with a human dataset. The chicken heart, naked mole rat spleen and green monkey lung were paired with the same tissue from Tabula Sapiens v1. For chicken retina, we transferred labels from a human retina dataset [8]. The dataset pairs were used to integrate datasets with SATURN using default parameters and author cell type annotations.

To perform label transfer, for a given dataset pair we first determine a set of common cell types. Next, we train a logistic classifier to predict cell type from the embeddings of the human dataset for the given model. The accuracy of the classifier is then assessed by applying it to the embeddings of the new species' cells. Since the classifier predicts cell types from the human dataset, its maximum possible accuracy is less than 1 as there are cells in the new species dataset which may not have a matched cell type label.

### Supplementary Note 6 Complex Dataset Benchmarking.

In order to evaluate the performance of UCE versus other foundation modeling approaches on complex neuronal datasets, we benchmark them on the Human Brain Cell Atlas [9]. To measure performance on a larger number of cell types, we benchmark "bio conservation" scores using the single cell integration benchmark (SCIB) [10] for the "cluster id" cell type resolution, which has 382 clusters. Because of the size of the dataset, which is almost $2.5$ million cells, we calculate scores using 10 randomly selected subsamples of $500,000$ cells. UCE outperforms scGPT and Geneformer on this benchmark, scoring an average bio conservation score of 0.611 (standard deviation 0.006), compared to scGPT (0.557, standard deviation 0.004) and Geneformer (0.434, standard deviation 0.009). UCE performance also matches or exceeds finetuned methods like scVI (Supplementary Table 10). These performance gains are also found in an additional dataset of $597,668$ cells taken from the developing mouse brain [11]. For the supercluster cell type column, which has $33$ unique cell types and the abca_subclass column, which has $282$ cell types, UCE (supercluster: 0.716, standard deviation 0.010, abca_subclass: 0.480, standard deviation 0.002) outperforms both scGPT (supercluster: 0.626, standard deviation 0.001, abca_subclass: 0.463, standard deviation 0.002) and Geneformer (supercluster: 0.487, standard deviation 0.002, abca_subclass: 0.420, standard deviation 0.002) in the zero-shot setting, calculated using 10 random samples of size $100,000$. Although UCE outperforms other methods in this zero-shot setting, for very fine cell type clusters, such as those found in the brain, traditional annotation approaches using marker gene expression and clustering should still be performed. These analyses can be supplemented by UCE outside of the label transfer setting, such as by performing initial clustering on the UCE space.

### Supplementary Note 7 Evaluation of Cell Type Organization in UCE space.

Computing a ground truth tree for cell type clusters is not trivial as the community does not hold a wide consensus on how cell types are defined and related to one another [12,13]. To our knowledge, Cell Ontology (CO) [4] is the most comprehensive attempt at a structured knowledge base for organizing cell types. We performed hierarchical clustering of all cell types from the Tabula Sapiens v2 Lung dataset based on the CO graph, using tree distances over the ontology. Here, we applied complete-linkage hierarchical clustering, meaning the distance between two clusters corresponded to the farthest tree distance between points in those clusters.

A challenge with using CO is that the granularity of certain cell types in the CO tree varies, potentially leading to longer tree distances even if the cell types are closely related. To address some of this uncertainty around true cell type labels, we used the clusterings obtained in the CO dendrogram at different depths of the dendrogram (normalized heights of 0.2, 0.4, 0.6). By visual inspection, these heights clearly corresponded to distinct levels of categorization (Supplementary Figure 12). At a height of 0.2, we observe clear separation of finer-grained cell type classes such as natural killer cells, smooth muscle cells, monocytes. At a height of 0.4, broader categories are identified such as all endothelial cells, secretory cells, pneumocytes. At the coarsest resolution of 0.6, major cell type categories are clearly distinguished. Overall, while we do not claim that CO is providing the 'ground truth' clustering, by leveraging its clustering at different levels, CO is an objective approximation that is useful for model benchmarking.

Next, we compared this tree to the clustering dendrogram inferred from the zero-shot embedding of previously unseen data from the Tabula Sapiens dataset (Figure 3a). We performed the same procedure for UCE, scGPT and Geneformer. We compare the dendrogram produced at each of the three levels of granularity of the CO tree with a range of different heights within the dendrograms produced by each embedding model (Supplementary Figure 13). This ensures that we do not choose a dendrogram height that unfairly biases any one model over the others. We measure overlap in clustering using the Adjusted Rand Index (ARI) between the clustering obtained through CO versus that obtained through one of the three model embeddings. The maximum value obtained for each model is recorded (Supplementary Figure 14).

The results show that at all depths, UCE embeddings are better at identifying CO clusters as compared to scGPT and Geneformer. The starkest contrast is visible at lower values of tree height (i.e. finer-grained cell type clustering) where it is harder to achieve good accuracy. Even at higher values of height, the UCE dendrogram shows more distinct separation of well-known cell type lineages such as immune cells, epithelial cell and endothelial cells as compared to scGPT and

Geneformer (Supplementary Figure 13).

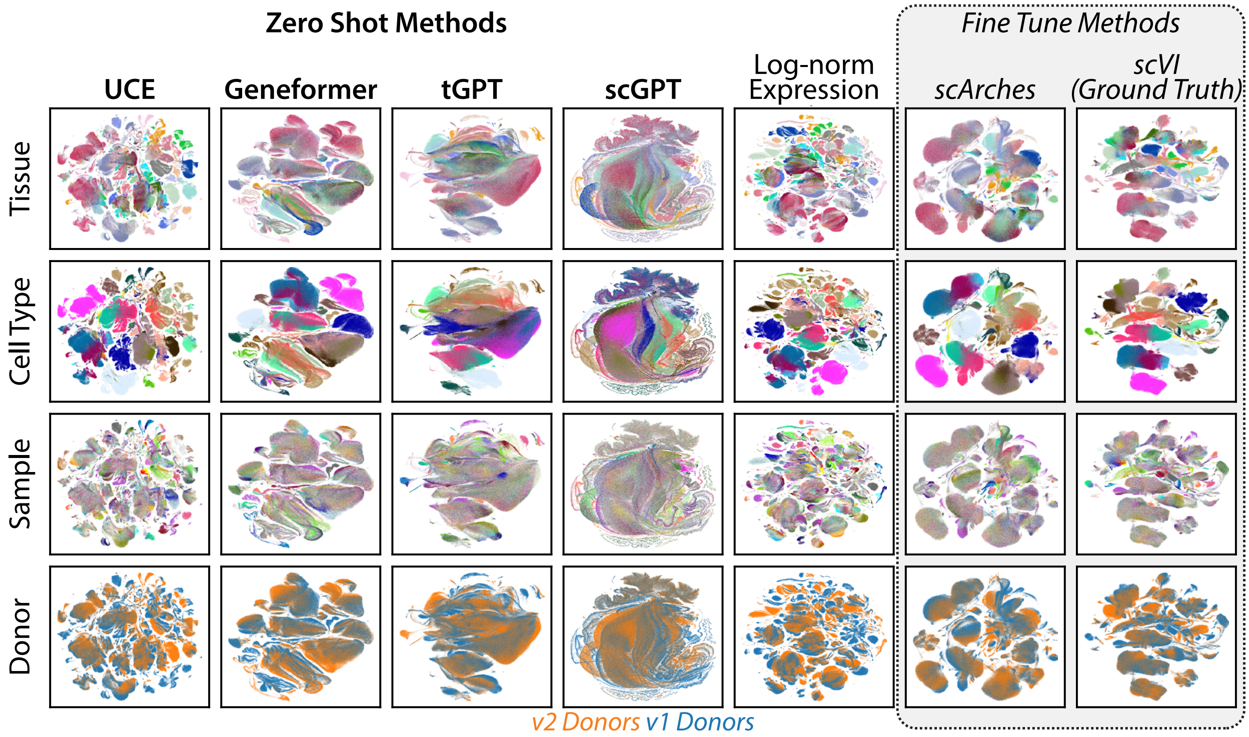


**Supplementary Figure 1: UCE outperforms existing methods for integrating Tabula Sapiens v2.** UMAP embeddings of Tabula Sapiens v2 and v1 donor colored by tissue, cell type, sample and donor group (v1 or v2). UCE's zero-shot embedding of Tabula Sapiens outperforms existing methods using comprehensive benchmarking metrics of cell embedding quality (Supplementary Table 1). For UCE, Geneformer, tGPT and log normalized expression, no additional gene selection was done. For scGPT and scArches, highly variable gene selection was performed on Tabula Sapiens v1, selecting the top 3000 highly variable genes, and the same gene set was used for generating embeddings for Tabula Sapiens v2. For scVI, 3000 highly variable genes were selected from cells from both Tabula Sapiens v1 and v2. scArches and scVI were trained using default settings, taking the sample as categorical covariate. For scArches, the cell type column ("cell_ontology_class") was used during SCANVI training.

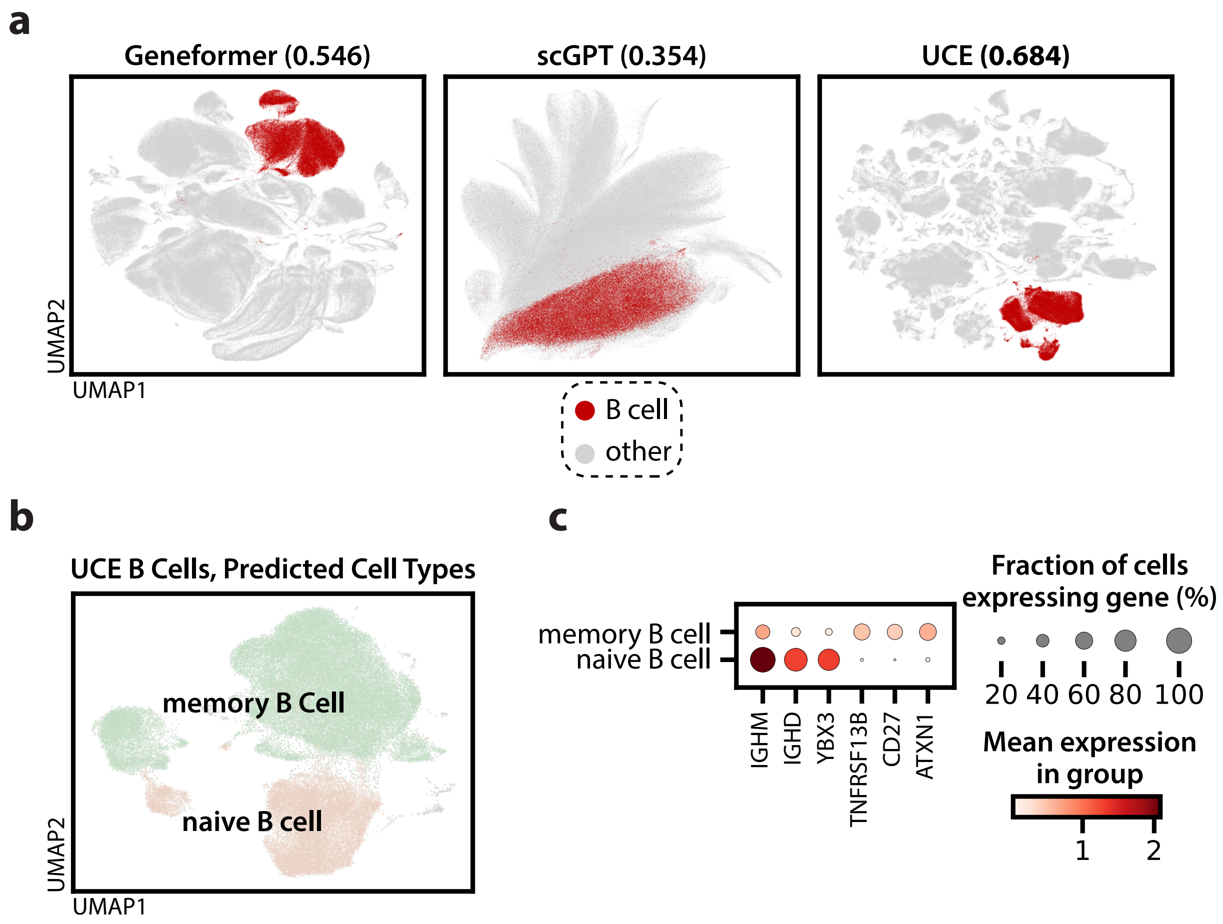


**Supplementary Figure 2: UCE embeddings recapture B cell identities in Tabula Sapiens v2.** **(a)** UMAP and B cell silhouette width scores of Tabula Sapiens v2 embeddings generated by Geneformer (left), scGPT (middle) and UCE (right), colored by cell type. The silhouette width score of B cells is 93% higher in UCE versus scGPT and 25% higher versus Geneformer. Cells annotated as B cells, colored in red, are distinctly clustered in the UCE space. In the Geneformer UMAP, B cells do form a large cluster, but a large number of other small clusters as well. In the scGPT UMAP, B cells are widely distributed. **(b)** UMAP of Tabula Sapiens v2 B cells, generated by UCE, colored by predicted cell type. A logistic classifier is trained on the UCE embeddings from the cross-tissue immune cell atlas [14], and then applied to predict cell types for Tabula Sapiens v2 B cells. Tabula Sapiens v2 B cells are indeed accurately classified as B cells of two categories, memory B cells and naive B cells. **(c)** Differential gene expression analysis between predicted memory B cells and predicted naive B cells. Known naive B cell markers *Ighm* and *Ighd* are preferentially expressed in the predicted naive B cells versus the predicted memory B cells[15]. Memory B cell markers *Tnfrsf13b* and *Cd27* are also preferentially expressed in the predicted memory B cells[14].

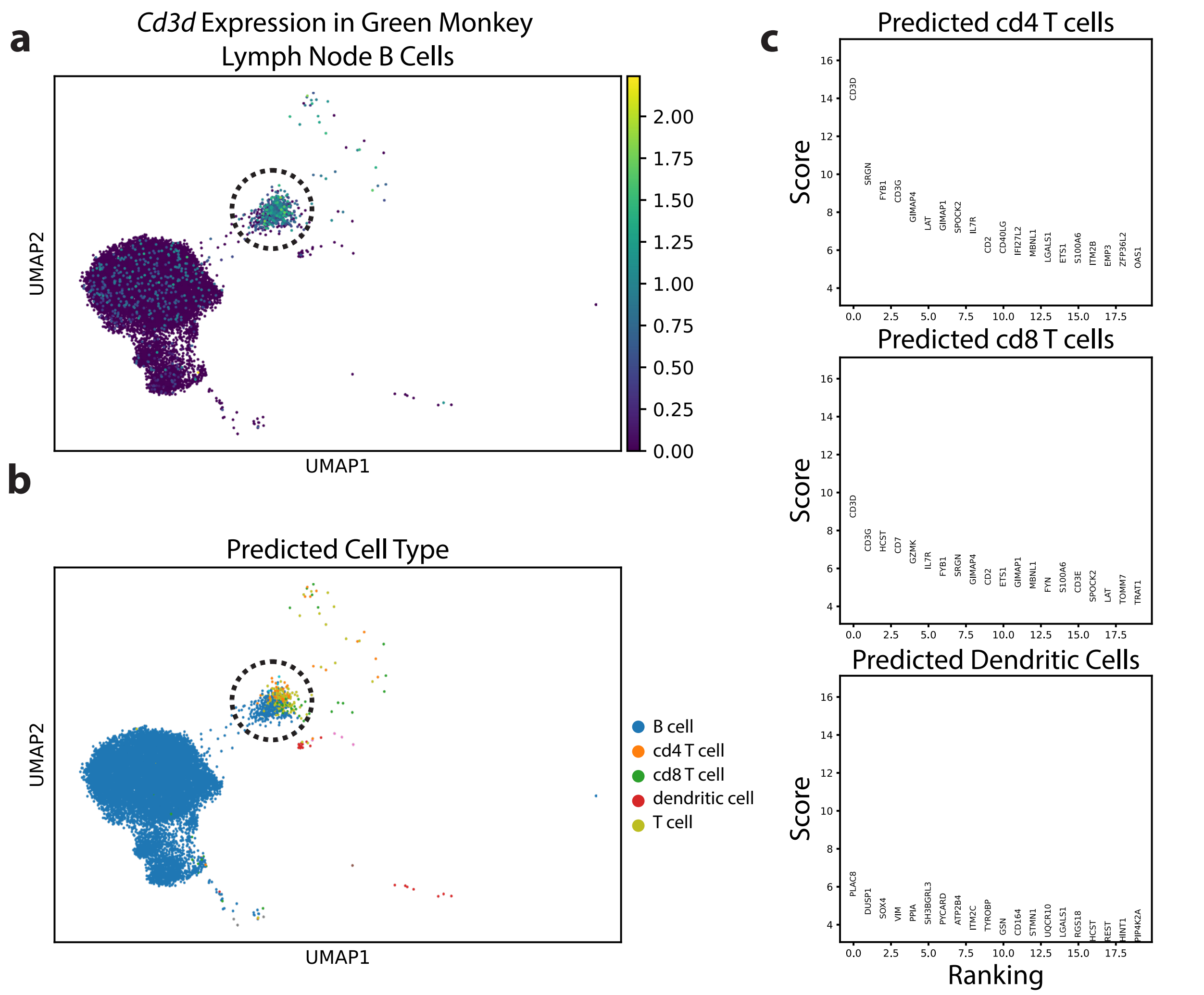


**Supplementary Figure 3: UCE captures *Cd3* expression in green monkey lymph node.** UMAP embeddings of cells originally labeled as B cells from green monkey lymph node, colored by **(a)** log-normalized expression of *Cd3d* and by **(b)** predicted human cell type. A population of cells, originally identified as B cells, forms a separate cluster, visually identifiable in the UMAP. This cluster preferentially expresses a T cell marker, *Cd3d*. A logistic classifier is trained to predict cell types from UCE embeddings for human lymph node cells from the IMA. The classifier is then applied to predict cell types for green monkey cells. **(c)** Differential expression for predicted T and dendritic cells versus predicted B cells, for cells originally annotated as B cells. Predicted T cells preferentially express *Cd3*, *Fyb1*, *Il7r*, *Hcst* and *Gzmk*, which are known markers for *Cd4* and *Cd8* T cells[16]. Supplementary RNA expression data from the human protein atlas confirms that these genes have higher RNA expression in T cells versus B cells [17]. Additionally, cells predicted to be dendritic cells preferentially express dendritic cell markers *Plac8*[18] and *Dusp1*[19].

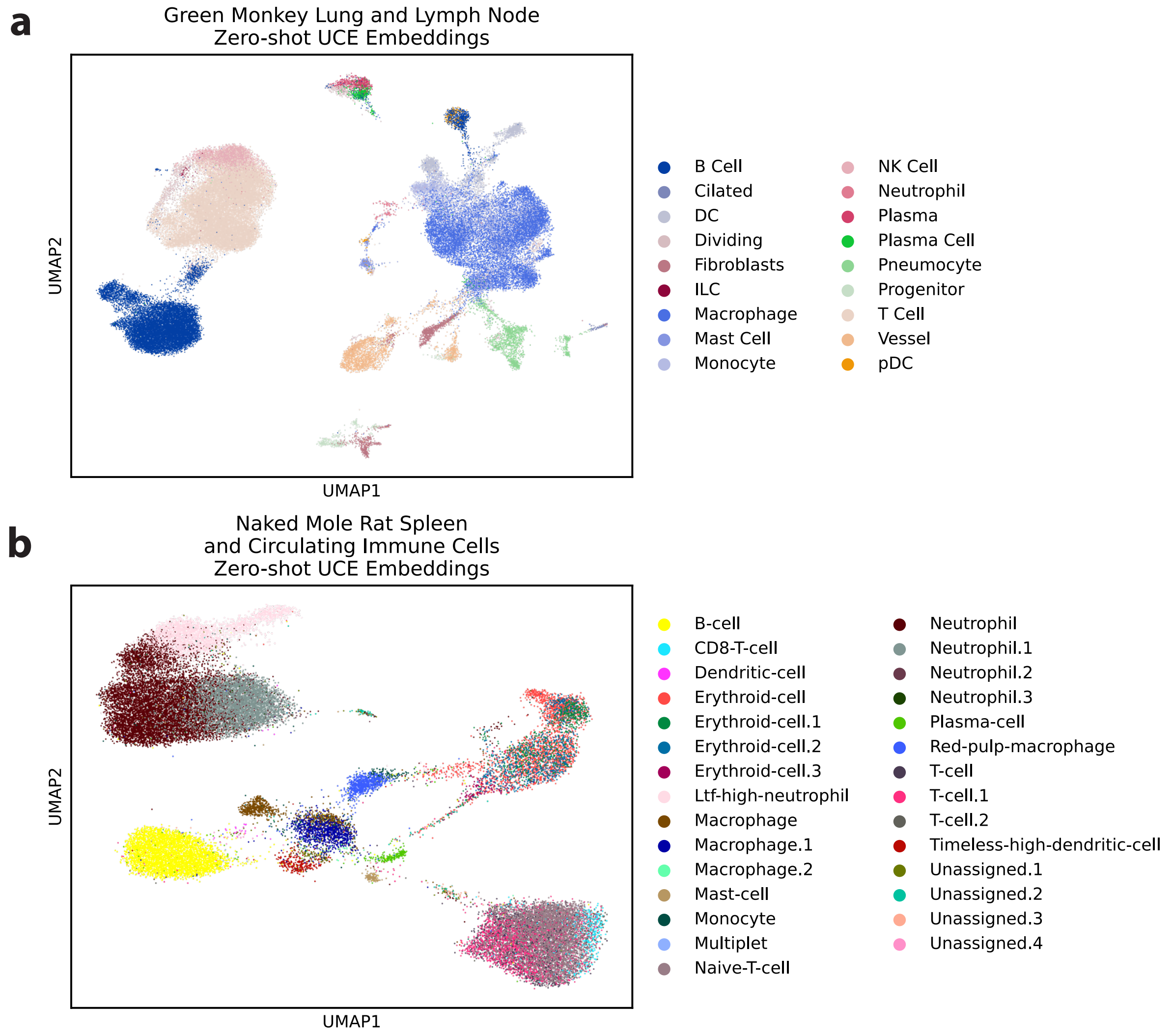


**Supplementary Figure 4: UCE can embed novel species that it was not trained on (green monkey, naked mole rat) in a zero-shot setting.** UMAP embeddings of green monkey lung and lymph node[20] (**a**), and naked mole rat spleen and circulating immune cells [21] (**b**) colored by cell type annotations. UCE is able to embed datasets from species it was not trained on in a zero-shot setting. UCE embeddings accurately recapture cell type information, and cell types in the new species map closely to their counterparts from other species in the IMA (Fig. 2d, Extended Data Table 1).

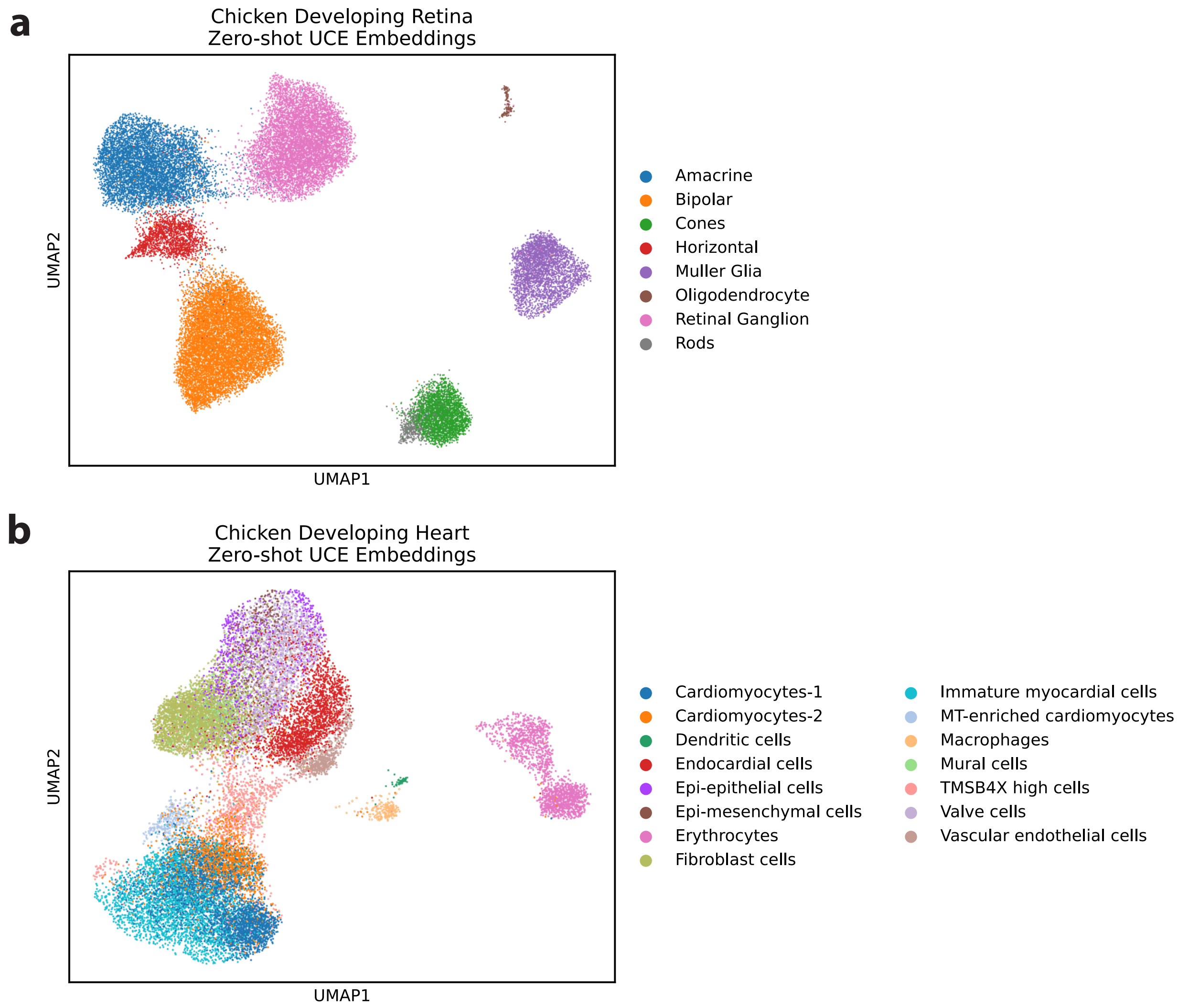


**Supplementary Figure 5: UCE can embed novel species that it was not trained on (chicken) in a zero-shot setting.** UMAP embeddings of chicken **(a)** retina [22] and **(b)** heart [23], colored by cell type annotations. UCE is able to embed datasets from species it was not trained on in a zero-shot setting. UCE embeddings accurately recapture cell type information, and cell types in the new species map closely to their counterparts from other species in the IMA (Extended Data Table 1).

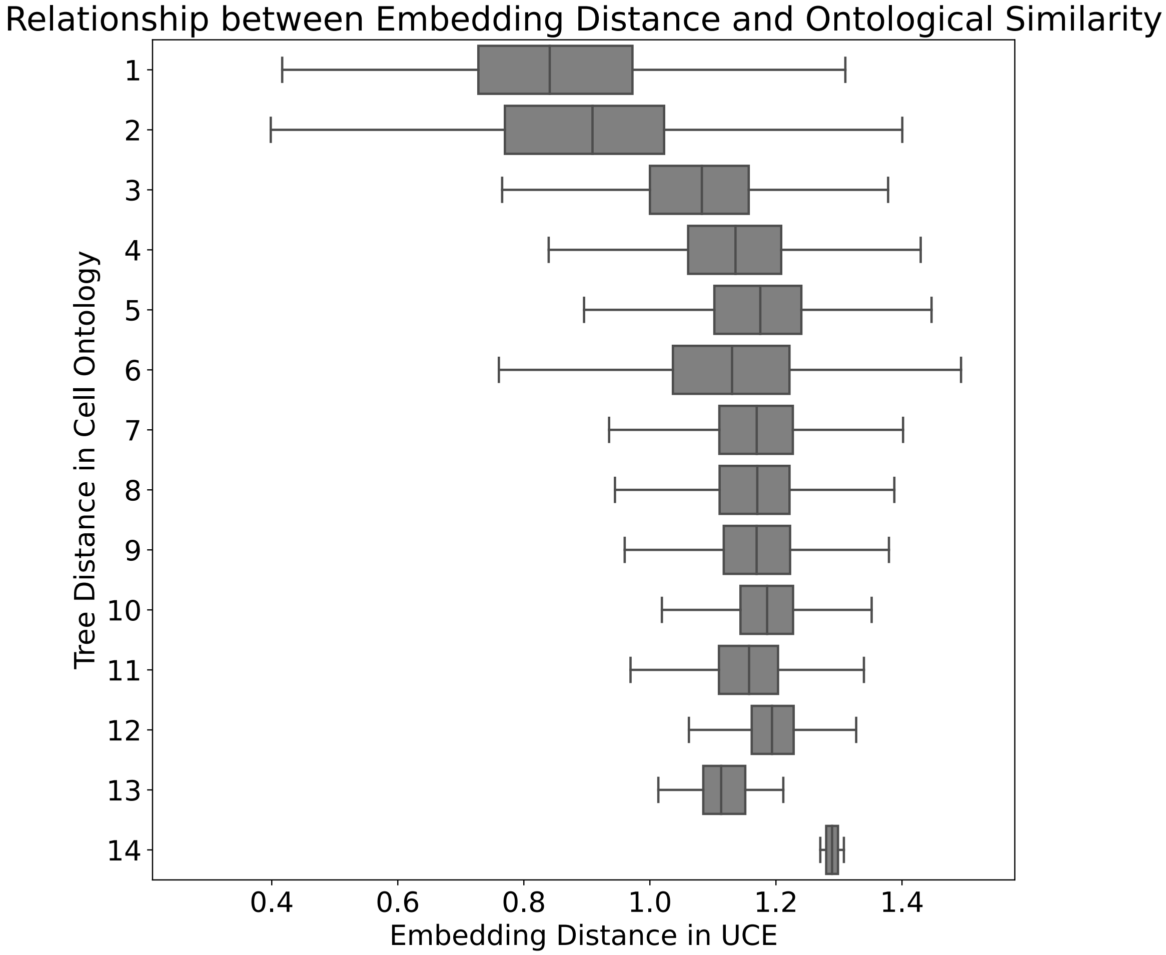


**Supplementary Figure 6: Relationship between Embedding Distance and Ontological Similarity** Relative organization of cell types in the embedding space compared to Cell Ontology. The x-axis depicts the Euclidean distance between all cell pairs across all tissues from Tabula Sapiens Consortium v2. The y-axis shows the corresponding tree distance between these cell types as found in the Cell Ontology.

**a**

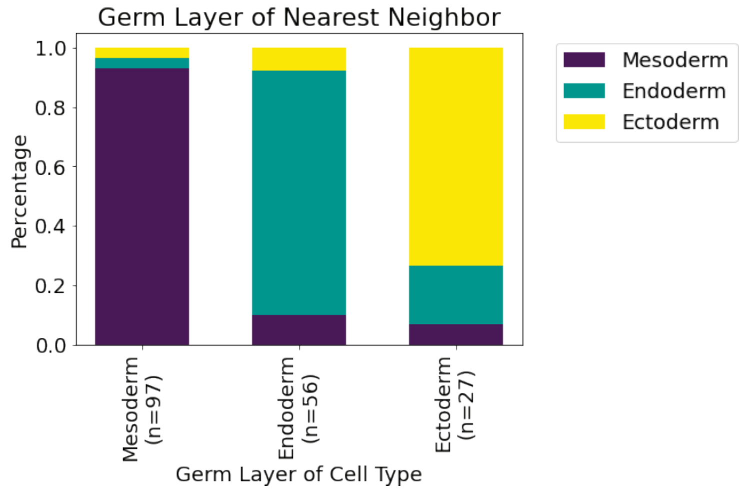


**b**

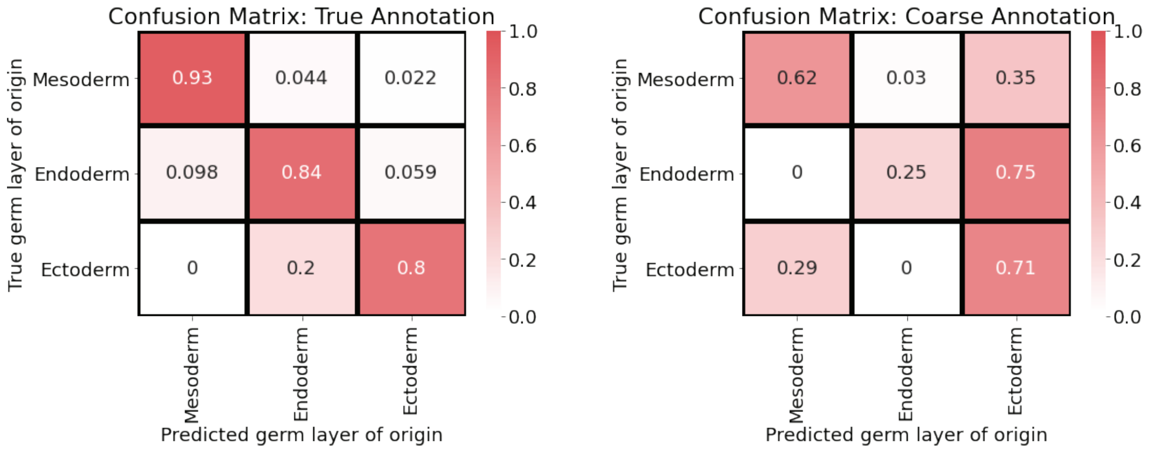


**Supplementary Figure 7: Cell types show organization in embedding space based on developmental lineage (a)** For each cell type centroid in the Tabula Sapiens v2 dataset, we determined the germ layer from which the closest neighboring cell type centroid originated. This information is presented in a stacked bar chart, summarizing the data for all cell types. On the x-axis, each bar represents a cell type associated with a different germ layer. The height of each segment within a stacked bar indicates the proportion of nearest centroids belonging to a specific germ layer. For instance, the leftmost bar shows that over 90% of the nearest centroids to mesoderm lineage cell types also originated from the mesoderm lineage. **(b)** We also present confusion matrices to demonstrate the accuracy of a classifier in predicting the germ layer of origin for cell types not included in the training set. The matrix on the left displays results from a classifier trained by excluding one cell type at a time. The matrix on the right, however, shows results from a model trained by excluding groups of related cell types. These related cell types were identified based on coarse annotations and were grouped for exclusion during the training process.

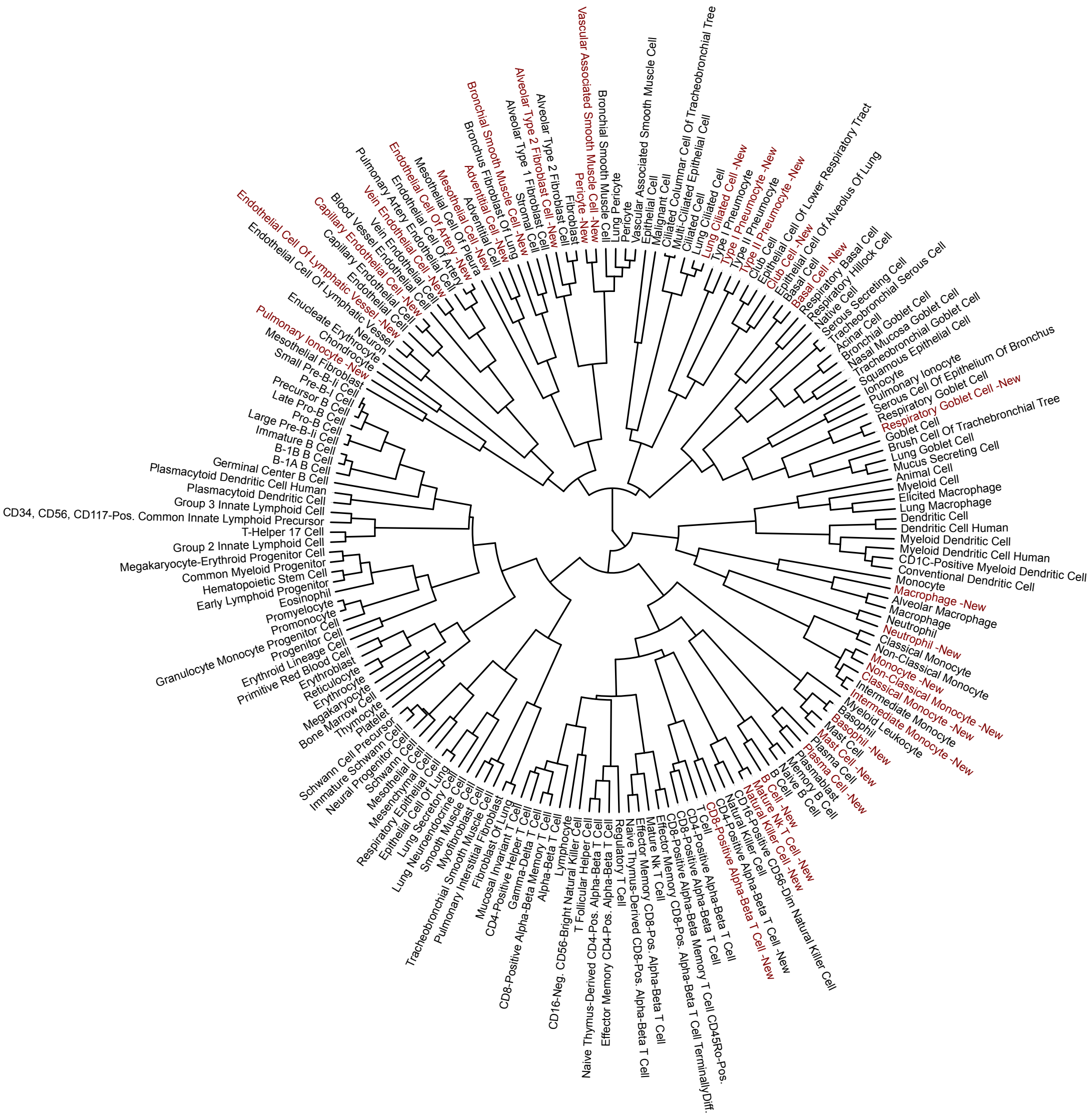


**Supplementary Figure 8: Mapping lung data from Tabula Sapiens v2 to multiple lung datasets from the Integrated Mega-scale Atlas (IMA)**. Red labels correspond to data from new donors (Tabula Sapiens v2), black are from IMA datasets. All cell type labels from multiple datasets are displayed as-is, with only small modifications to abbreviate longer labels.

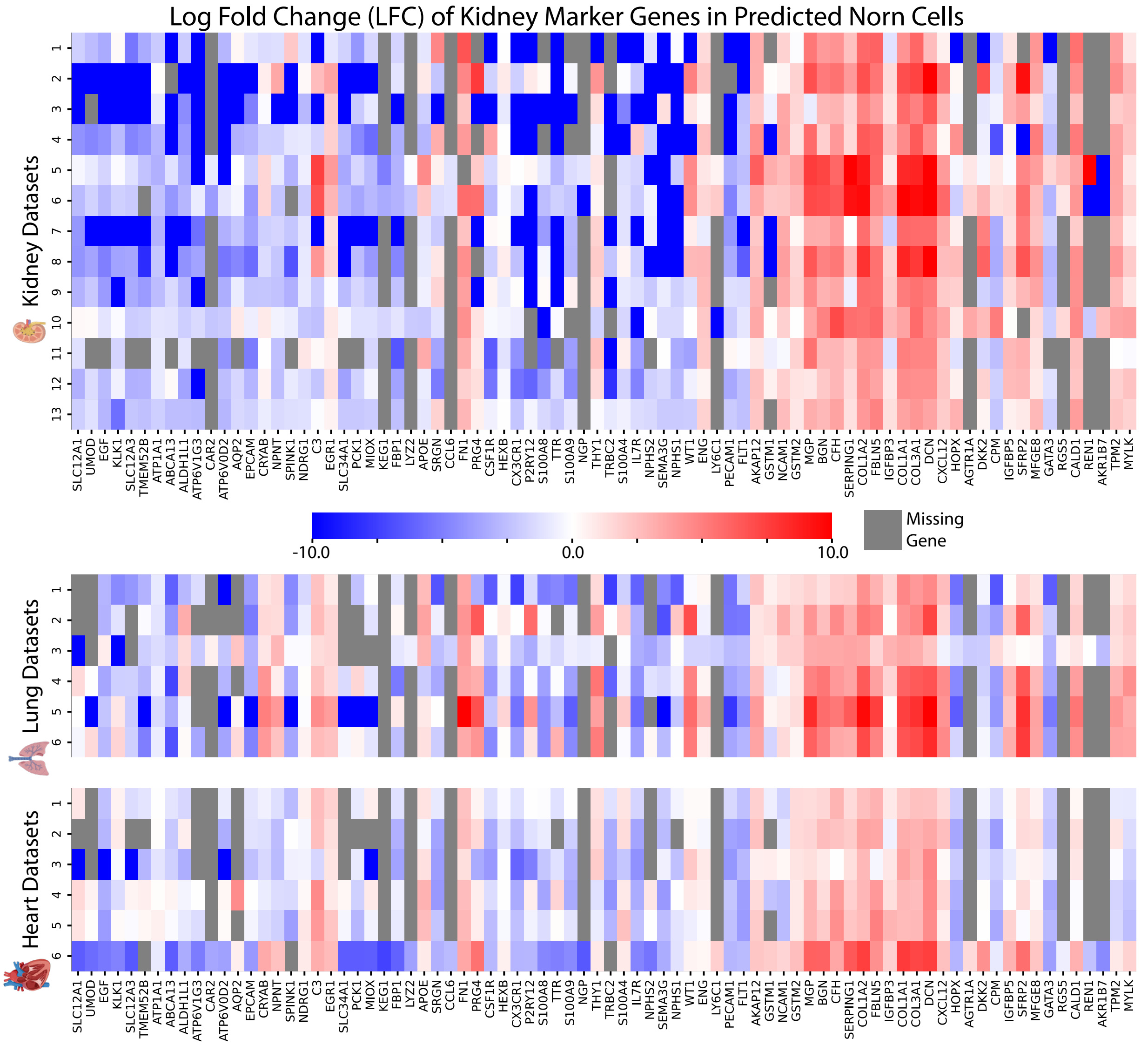


**Supplementary Figure 9: Predicted Norn cells express canonical Norn Markers.** Identification of Norn cells and Norn-like cells across tissues. A logistic classifier is trained to predict Norn cells from universal cell embeddings for mouse kidney cells[24], and is applied to kidney datasets (top) and datasets from lung and heart (bottom). The log fold change of kidney cell type marker genes is visualized between cells predicted to be Norn cells and the remaining cells within each dataset. Cells which are predicted to be Norn-like indeed have high expression of Norn marker genes like *Col1a1* and *Dcn*[24] (Extended Data Table 3).

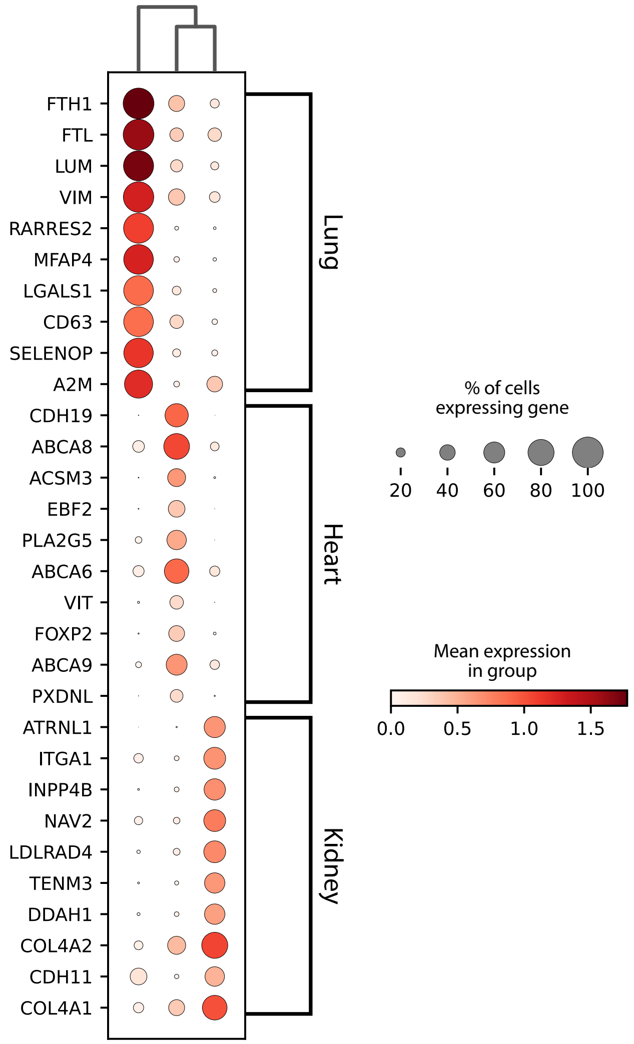


**Supplementary Figure 10: Differential Expression Between Norn-like cells Across Tissues.** Results of differential expression analysis between predicted Norn-like cells in datasets from three tissues, lung[25], heart[26] and kidney[27]. Predicted Norn-like cells in the lung preferentially express ferritin components *Fth1* and *Ftl* [28,29]. Norn-like cells in the heart preferentially express *Cdh19*, which is broadly expressed in heart tissues[17,28,29].

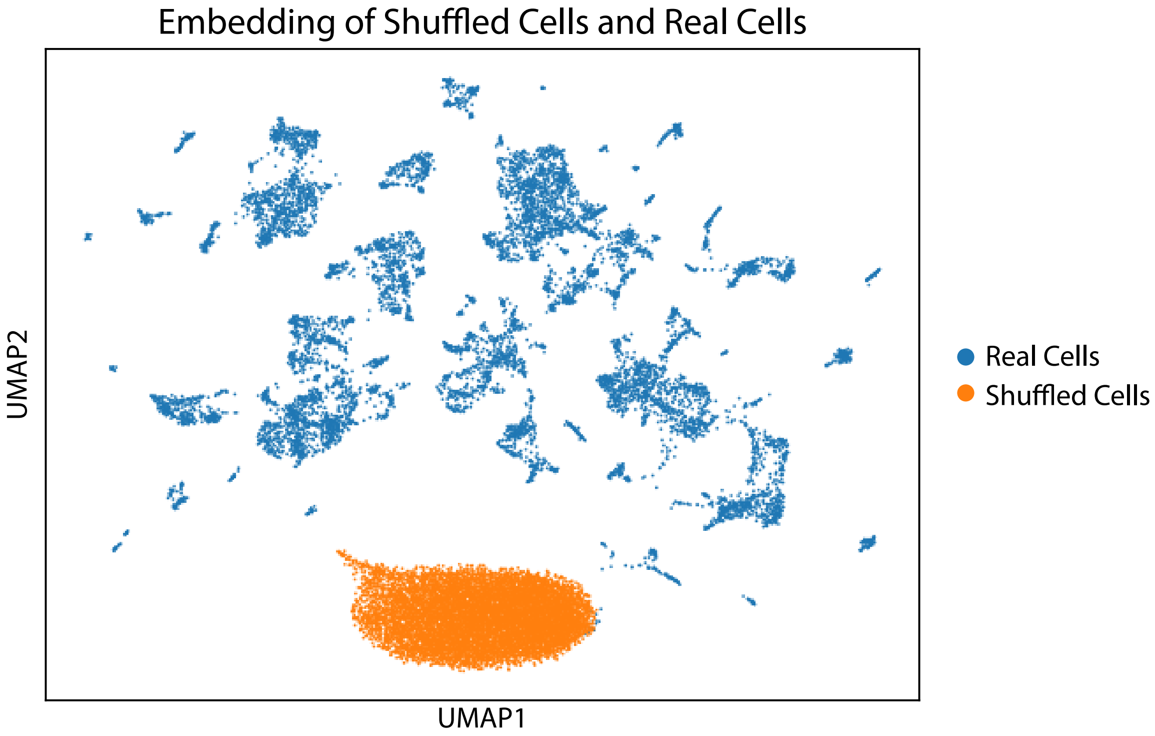


**Supplementary Figure 11: Embedding of Fake Cells with UCE.** UMAP calculated from Universal Cell Embeddings of $10,000$ cells randomly sampled from Tabula Sapiens v1 and v2, colored in blue, and shuffled versions of those cells in orange. To shuffle the cells, the count values of all $10,000$ cells are randomly shuffled without respect to row (cell) or column (gene) ordering. The new, shuffled cells are then embedded using UCE. Embeddings of the shuffled cells are highly out of distribution and homogeneous. A one-nearest neighbor classifier trained on the real $10,000$ cells predicts that for $8,822$ of the fake cells, the nearest neighbor is the same exact real cell, a spermatid cell.

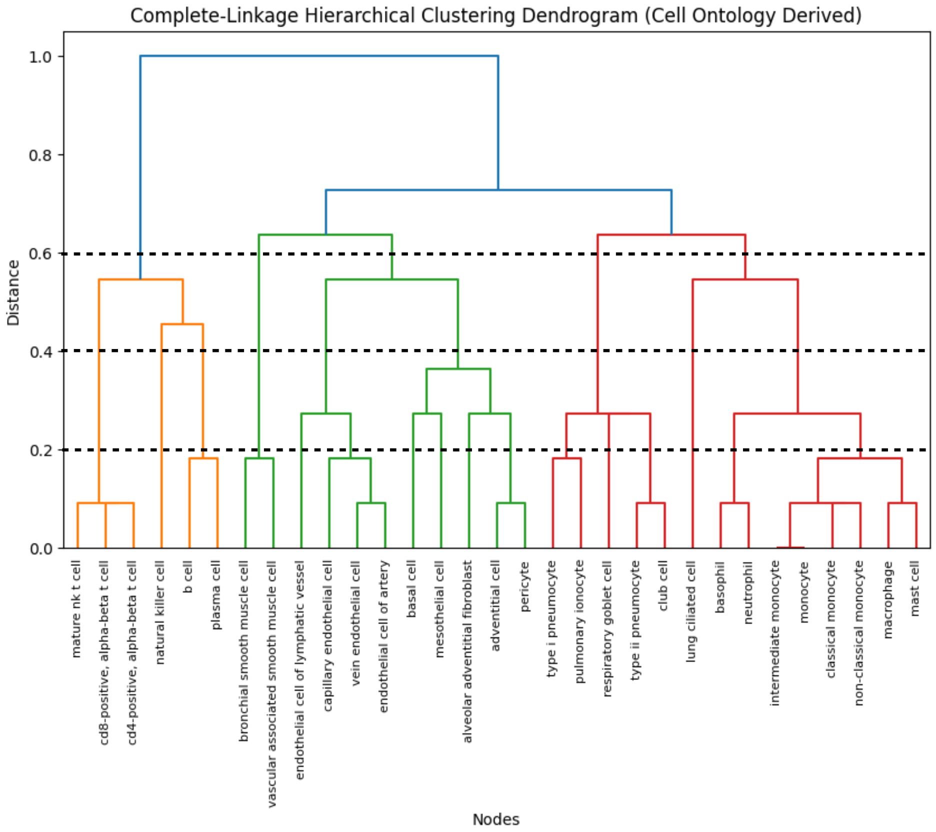


**Supplementary Figure 12: Using Cell Ontology to derive Reference Lung Cell Type Organization** Cell type clustering derived from complete-linkage hierarchical clustering of lung cell types in Cell Ontology. Lung cell types were obtained from the Tabula Sapiens v2 dataset. Dotted lines correspond to different heights at which the dendrogram was cut to obtain clusterings of varying resolutions.

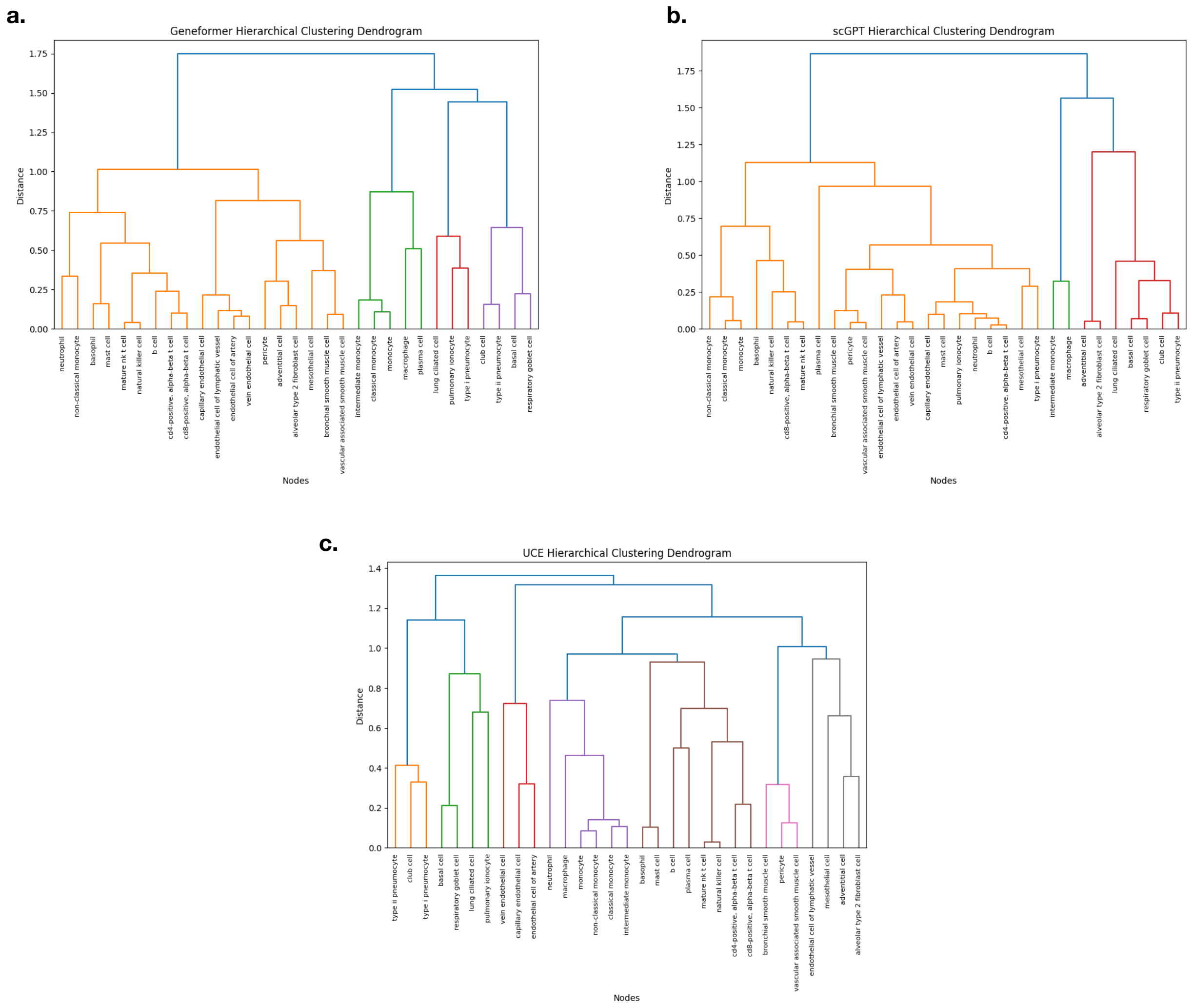


**Supplementary Figure 13: Dendrograms of Cell Type Organization created by different cell embedding methods** Dendrogram of cell type clustering generated using (a) Geneformer (b) scGPT (c) UCE. Dendrogram for each model was generated using that model's embedding of new, previously unseen data from Tabula Sapiens v2.

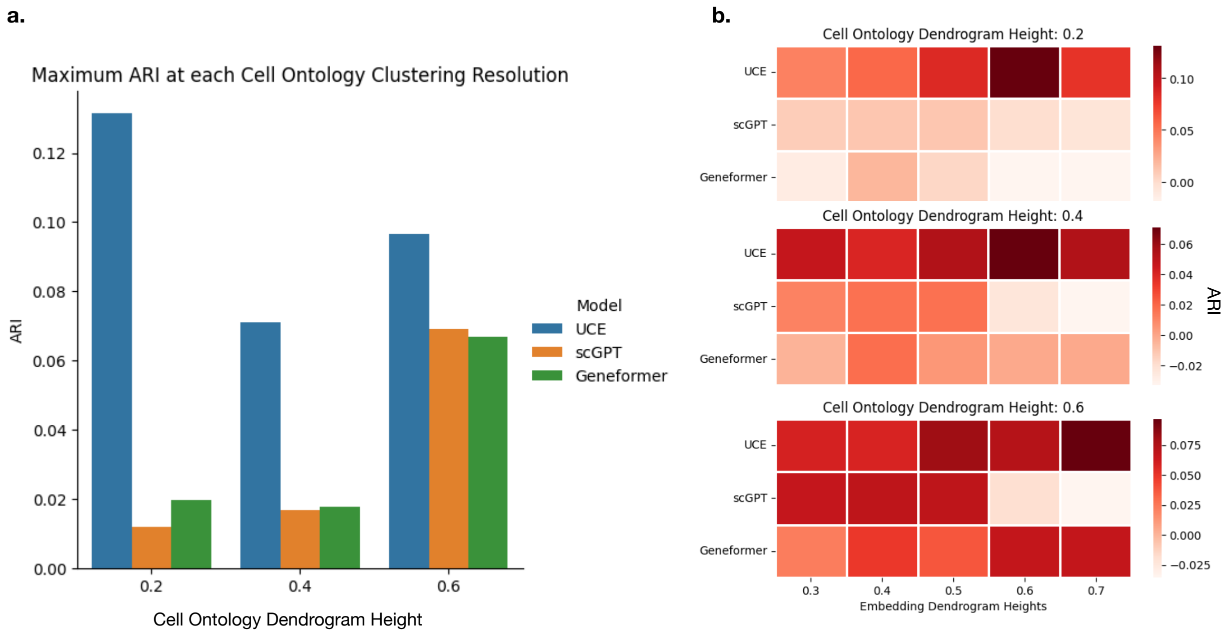


**Supplementary Figure 14: Evaluation of Cell Type Organization in UCE space** The embedding space for each model was generated for new, previously unseen data from Tabula Sapiens v2. (a) Adjusted Rand Index computed at each clustering resolution of the Cell Ontology dendrogram (Supplementary Figure 12) (0.2, 0.4, 0.6) compared to clusterings produced by three different cell embedding methods applied zero-shot to the Tabula Sapiens dataset. (b) At each clustering resolution of the Cell Ontology dendrogram, each embedding model was evaluated at 5 clustering resolutions (0.3, 0.4, 0.5, 0.6, 0.7). The maximum value for each model was used for (a).

Large Intestine (Tabula Sapiens v2) UCE Embedding

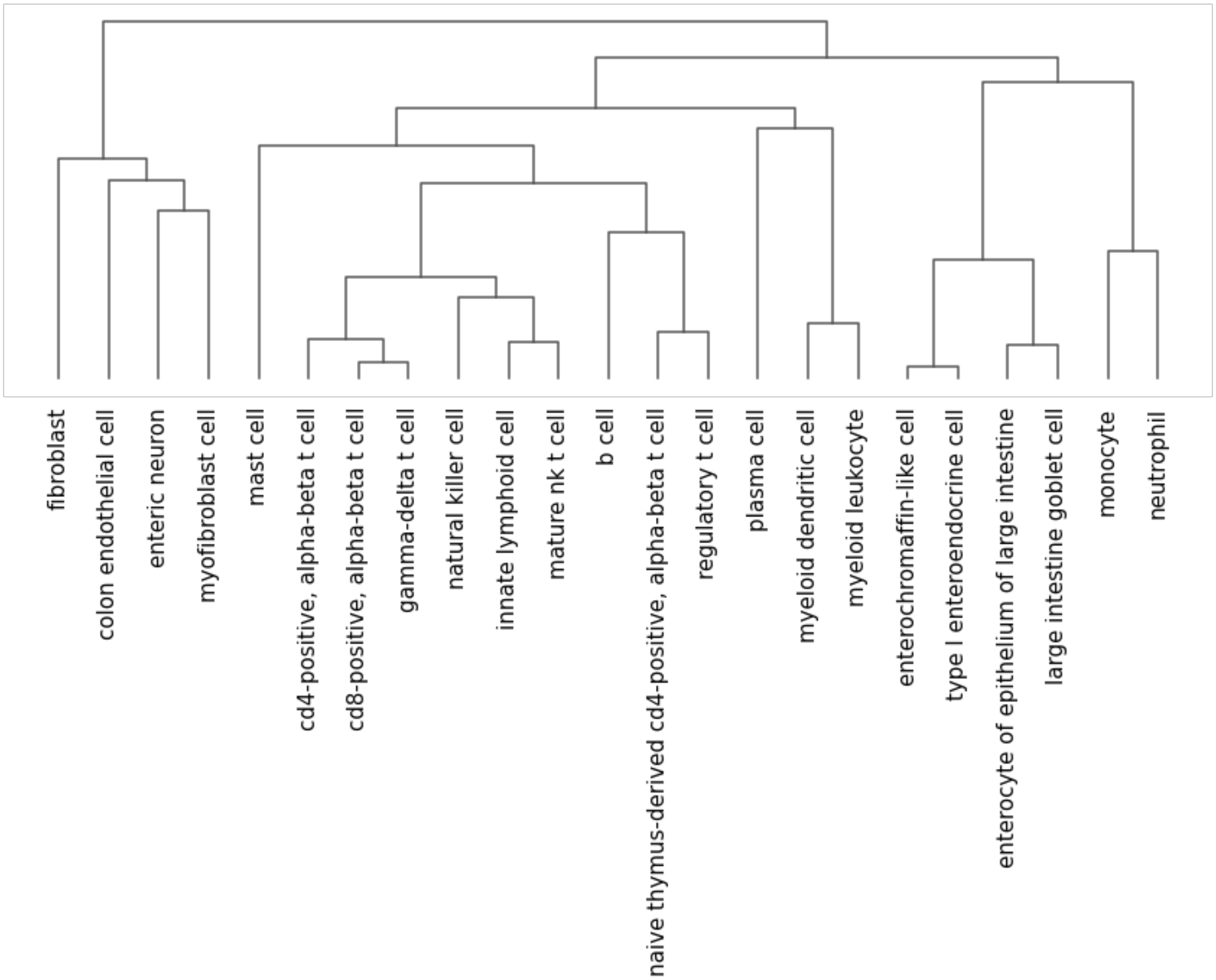


**Supplementary Figure 15: Hierarchical clustering of cell types in the Large Intestine in the UCE space identifies ontological relationships** The UCE space was generated for new, previously unseen data from Tabula Sapiens v2. We observe that T-cell subtypes and natural killer cells are separated from other leukocytes such as dendritic cells and plasma cells. Fibroblasts are separated from leukocytes and cluster distinctly from all leukocytes. There are also instances where UCE does not correctly disentangle finer cell lineages: some T-cell subtypes cluster distinctly from others and are in fact closer to B-cells.

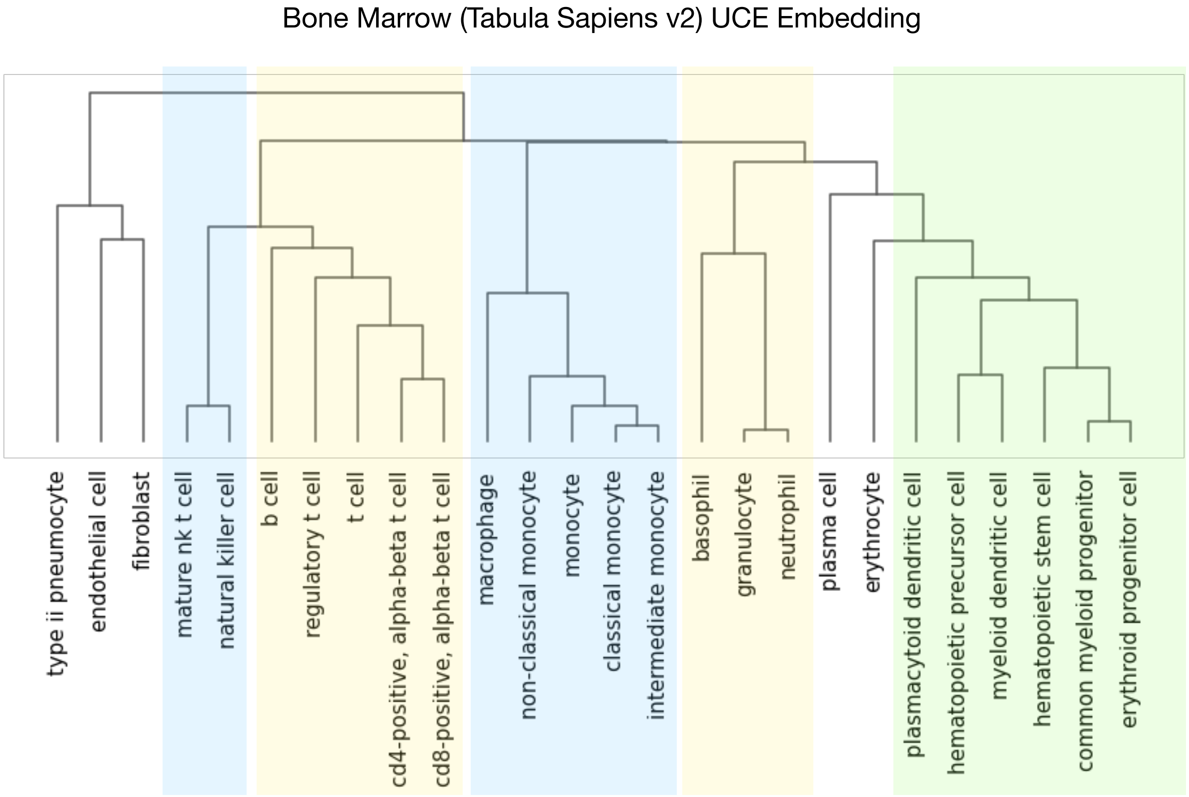


**Supplementary Figure 16: Hierarchical clustering of cell types in the Bone Marrow in the UCE space identifies developmental relationships** The UCE space was generated for new, previously unseen data from Tabula Sapiens v2. While early erythroid and myeloid progenitors cluster closer to each other, more advanced stages within distinct lineages, such as the lymphoid lineage (T-cells, B-cells and NK cells) and the myeloid lineage (macrophages, monocytes, granulocytes), cluster distinctly from one another. Within individual lineages, we observe macrophages and monocytes clustering distinctly from granulocytes such as neutrophils and basophils.

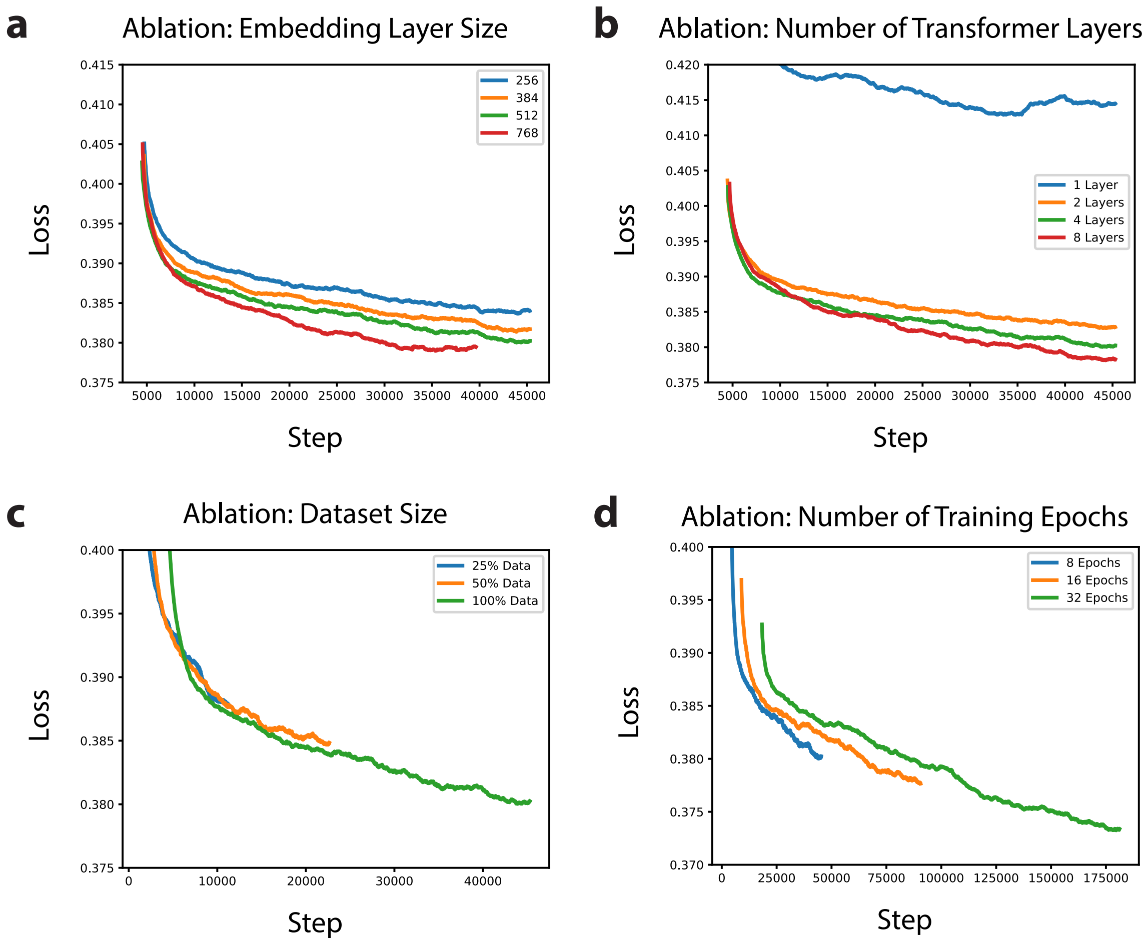


**Supplementary Figure 17: Model and Dataset Size Ablations** Loss versus step is plotted for different groups of model ablations. Increasing model size, either by increasing the transformer layer embedding dimension (a) or number of transformer layers (b) reduced loss. Training of the 768 embedding dimension model diverged during the final epoch, so results are shown for the first 7 epochs. Model performance improves as training dataset size (c) or training length (d) increases. Loss is plotted as the rolling average of the previous 1000 steps. Details of ablations are further described in Supplementary Note 4.

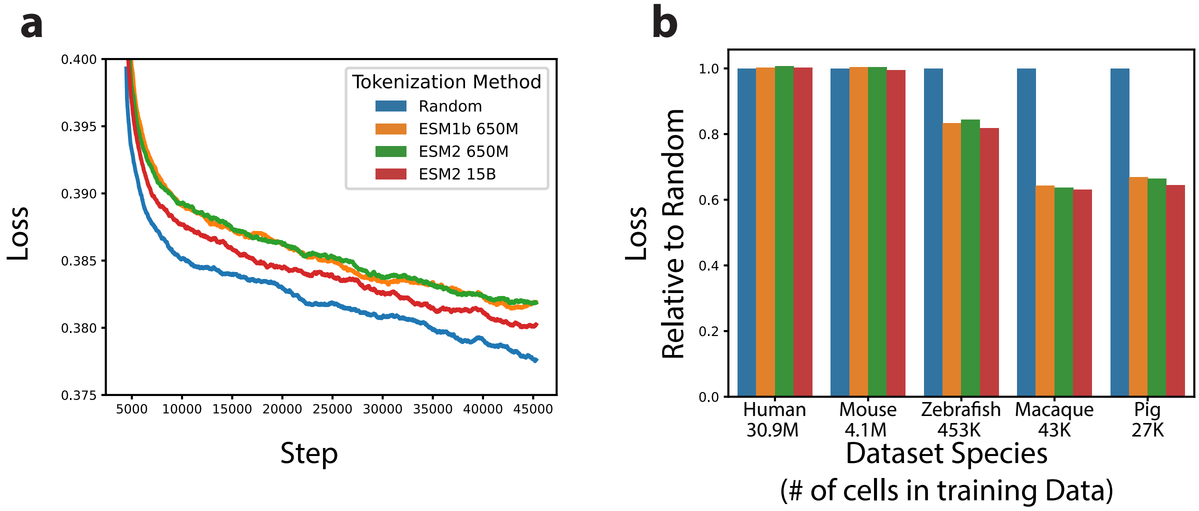


**Supplementary Figure 18: Protein Embedding Tokenization Ablations** (a) Loss versus step is plotted for different model ablations using either randomly initialized gene embeddings or protein language model embeddings. (b) Loss for the same models is evaluated on 5 different holdout datasets [30–32] from different species included in the training data. Loss values are scaled relative to the loss of the random gene token model. Protein language model tokenization improves performance on all non-human species. ESM2 15B outperforms ESM1b 650M and ESM2 650M on all species. Details of ablations are further described in Supplementary Note 4.

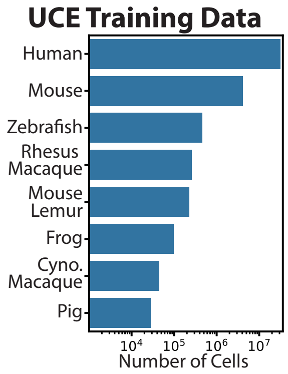


**Supplementary Figure 19: Training Data Species Distribution** Number of cells per each species in the training data.

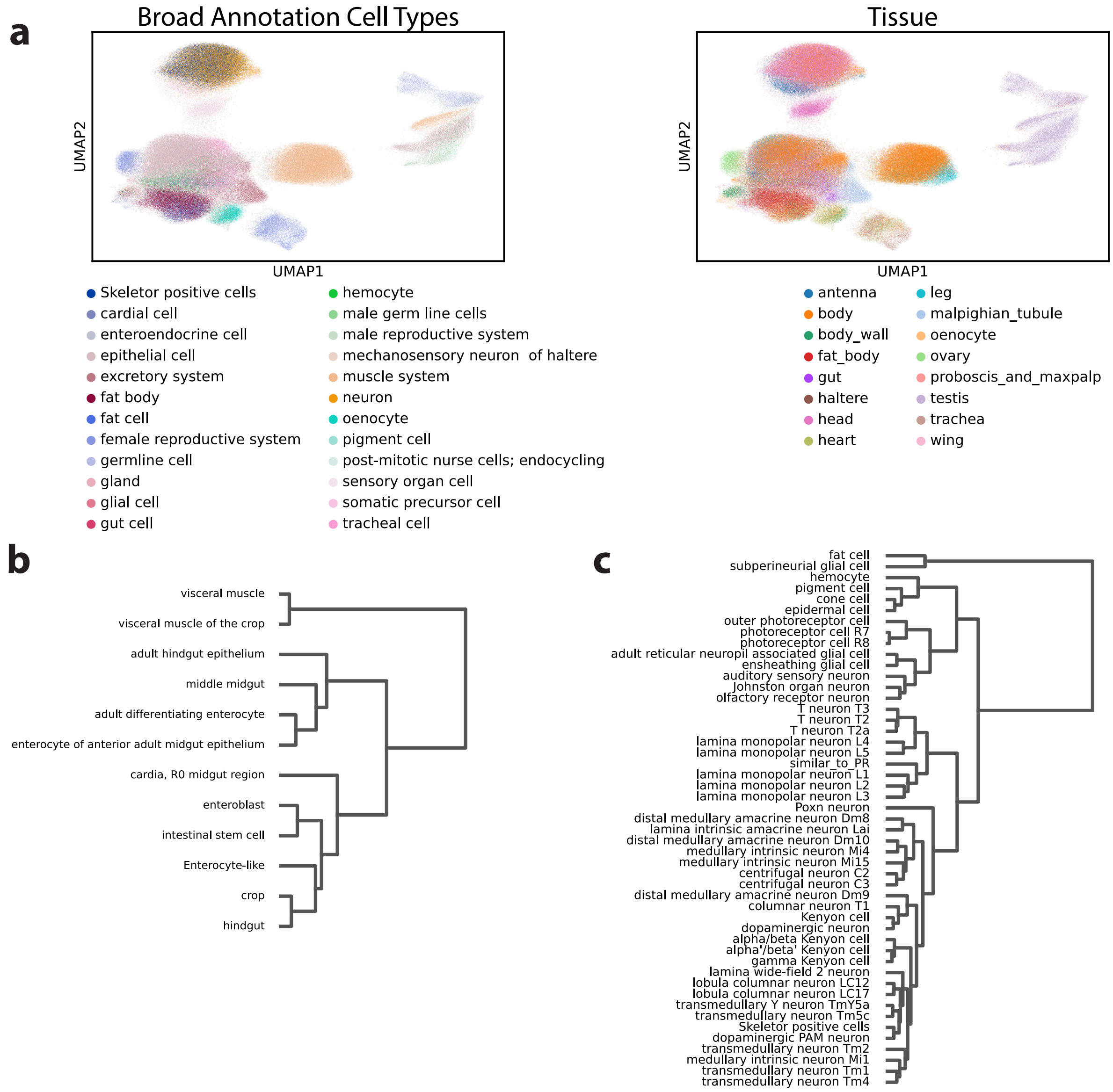


**Supplementary Figure 20: Zero Shot Fly Cell Atlas Embedding** Zero shot embeddings for the fly cell atlas [33] are generated with UCE. (a) UMAP visualization of the UCE embeddings colored by broad annotation cell types (left) and tissue (right). A logistic regression classifier trained on the UCE embeddings of the dataset can classify its cell types with 80.6% accuracy (10 fold cross validation). A random classifier in comparison, achieves only 22% accuracy. Cells with the broad annotation 'unknown' and 'artifact' are removed. Hierarchical dendrograms, generated based on distance between consensus community cell types in the zero shot embedding space, for different fly tissues, (b) gut (c) head. Cell types with 100 or more cells are retained.

| Embedding Method | Isolated labels | KMeans NMI | KMeans ARI | Silhouette label | cLISI | Silhouette batch | iLISI | KBET | Graph connectivity | Batch correction | Bio conservation | Total Score |
|---|---|---|---|---|---|---|---|---|---|---|---|---|
| **Zero Shot Methods** | | | | | | | | | | | | |
| UCE | **0.648** (0.0046) | **0.703** (0.002) | **0.229** (0.0124) | 0.537 (0.0003) | **1** (0.0) | **0.878** (0.0012) | 0.007 (0.0) | 0.412 (0.0067) | 0.694 (0.0032) | **0.498** (0.0017) | **0.623** (0.003) | **0.573** (0.0014) |
| scGPT | 0.452 | 0.29 | 0.039 | 0.404 | 0.994 | 0.816 | **0.036** | 0.295 | 0.349 | 0.374 | 0.436 | 0.411 |
| tGPT | 0.555 | 0.497 | 0.111 | 0.45 | 0.999 | 0.876 | 0.007 | 0.302 | 0.598 | 0.446 | 0.522 | 0.492 |
| Geneformer | 0.531 | 0.554 | 0.143 | 0.454 | 0.999 | 0.847 | 0.007 | 0.312 | 0.644 | 0.452 | 0.536 | 0.503 |
| **Raw Data** | | | | | | | | | | | | |
| Log-Normalized Expression, PCA | 0.614 | 0.673 | 0.189 | 0.519 | **1** | 0.828 | 0.003 | 0.332 | 0.746 | 0.477 | 0.599 | 0.55 |
| **Fine Tuned Methods** | | | | | | | | | | | | |
| scVI | 0.605 (0.0085) | 0.67 (0.0016) | 0.195 (0.0036) | 0.51 (0.0031) | 0.999 (0.0) | 0.823 (0.0018) | 0.016 (0.0002) | **0.419** (0.0079) | 0.695 (0.0054) | 0.488 (0.0022) | 0.596 (0.0019) | 0.553 (0.0013) |
| scArches | 0.623 (0.0179) | 0.679 (0.0054) | 0.224 (0.0128) | **0.544** (0.0049) | 0.999 (0.0001) | 0.819 (0.0076) | 0.017 (0.0007) | 0.409 (0.0169) | **0.724** (0.009) | 0.492 (0.0078) | 0.614 (0.0058) | 0.565 (0.005) |

**Supplementary Table 1: UCE Performance on single-cell Integration Benchmark** Model performance on Tabula Sapiens v2 evaluated against other methods in the zero-shot setting. Two fine-tuned methods were also included as a baseline for assessing performance. Metrics are divided into those that assess cell type alignment performance and those that measure effectiveness of batch effect correction [10]. Overall score takes the weighted average over cell type matching score and batch correction score $(0.6 * \text{Avg. Bio}) + (0.4 * \text{Avg. Batch})$. For certain models results have a small amount of variance, either because the model is fine tuned model (scVI, scArches), or because the model includes sampling (UCE). For these models, the dataset is shuffled using 10 different random seeds, and then the method is evaluated (used to create embeddings) and scored. For the fine tuned models, a new model was trained on the shuffled data for each seed.

| Hyperparameter | Value |
|---|---|
| Total Number of Parameters | 674,745,857 |
| Transformer Layers Number of Parameters | 649,355,520 |
| Number of Transformer Layers $(n_{lay})$ | 33 layers |
| Transformer Layer Embedding Size, $(d_{emb})$ | 1280 |
| Transformer Layer Hidden Dimension $(d_{hid})$ | 5120 |
| Number of Transformer Heads $(n_{head})$ | 20 heads |
| Dropout | 0.05 |
| Transformer Layer Activation Function | ReLU |
| Transformer Layer Normalization | LayerNorm |
| Number of Genes Sampled per cell ($G_i^s$) | 1024 genes |
| Total Training Epochs | 8 epochs |
| Warmup Epochs | 1 epoch |
| GPU type | NVIDIA A100-SXM4-80GB |
| Number of Nodes | 3 machines |
| Total Number of GPUs | 24 GPUs |
| Per GPU Batch Size | 6 cells |
| Gradient Accumulation Steps | 4 steps |
| Effective Batch Size | 576 cells |
| CLS Token Decoder MLP Layer Dimensions (Transformer) | 5120, 1024, 1280, 1280 |
| Gene Token Embedding MLP Layer Dimensions (Input) | 5120, 1280 |
| Gene Token Embedding MLP Layer Dimensions (Loss) | 5120, 1280 |
| Binary Decoder MLP Layer Dimensions (Loss) | 2560, 2048, 512, 128, 1 |
| MLP Layer Activation Function | GELU |
| MLP Layer Normalization | LayerNorm |
| Expressed Genes per Cell $(\|G_i^{L+}\|)$ | 512 genes |
| Non-expressed Genes per Cell $(\|G_i^{L-}\|)$ | 512 genes |
| Gene Masking $(r_{mask})$ | 20% |
| Total Training Time | 43.5 days |

**Supplementary Table 2: UCE Model Hyperparameters** Model size was chosen based on available computational resources. Model size was chosen to be the same as another biological foundation model, ESM1b[1].

| Change to Ablation Base Model | Loss |
|---|---|
| Base Ablation Model: Positional Encoding | **0.3827** |
| Remove Positional Encoding | 0.3835 |

**Supplementary Table 3: Model Ablation: Chromosome Positional Encoding.** The 4-layer 22.1 million parameter base ablation model (Supplementary Note 4) is compared to an identical model, trained without positional encodings, to measure the impact of chromosome and genome location information on model performance. Performance is evaluated on the holdout dataset Tabula Sapiens v2.

| Change to Ablation Base Model | Loss |
|---|---|
| Base Ablation Model: Mask Rate 20% | **0.3827** |
| Decrease mask rate to 10% | 0.3833 |
| Increase mask rate to 40% | 0.3870 |

**Supplementary Table 4: Model Ablation: Gene Masking Rate.** The 4-layer 22.1 million parameter base ablation model (Supplementary Note 4) is compared to identical models, trained with either a decreased mask rate of 10%, or an increased mask rate of 40%. Performance is evaluated on the holdout dataset Tabula Sapiens v2.

| Embedding Method | Isolated labels | KMeans NMI | KMeans ARI | Silhouette label | cLISI | Silhouette batch | iLISI | KBET | Graph connectivity | Batch correction | Bio conservation | Total Score |
|---|---|---|---|---|---|---|---|---|---|---|---|---|
| UCE | 0.564 (0.009) | **0.469** (0.010) | **0.123** (0.013) | **0.611** (0.023) | 1 (0.0) | **0.906** (0.005) | 0.039 (0.000) | 0.581 (0.010) | **0.849** (0.014) | 0.594 (0.005) | **0.553** (0.005) | **0.569** (0.003) |
| scVI | 0.594 (0.009) | 0.37 (0.005) | 0.071 (0.003) | 0.575 (0.008) | 1 (0.) | 0.85 (0.003) | 0.079 (0.002) | **0.661** (0.025 | 0.838 (0.007) | **0.607** (0.008) | 0.522 (0.003) | 0.556 (0.004) |
| scArches | 0.593 (0.017) | 0.387 (0.016) | 0.079 (0.005) | 0.588 (0.022) | 1 (0.0) | 0.853 (0.004) | 0.08 (0.016) | 0.617 (0.049) | 0.833 (0.016) | 0.596 (0.017) | 0.529 (0.008) | 0.556 (0.010) |
| PCA | **0.613** | 0.451 | 0.089 | 0.555 | 1 | 0.857 | 0.003 | 0.382 | 0.825 | 0.517 | 0.542 | 0.532 |
| Geneformer | 0.559 | 0.36 | 0.09 | 0.479 | 1 | 0.873 | 0.025 | 0.357 | 0.749 | 0.501 | 0.498 | 0.499 |
| tGPT | 0.562 | 0.291 | 0.052 | 0.462 | 1 | 0.873 | 0.014 | 0.231 | 0.659 | 0.444 | 0.474 | 0.462 |
| scGPT | 0.45 | 0.069 | 0.01 | 0.378 | 0.997 | 0.838 | **0.116** | 0.395 | 0.498 | 0.461 | 0.381 | 0.413 |

**Supplementary Table 5: Performance on Tabula Sapiens v2 Ovary Tissue** Model performance on Tabula Sapiens v2 Ovary tissue evaluated against other methods in the zero-shot setting. The ovary tissue contains cells measured with 10x-primev3 and with Smart-seq3. Metrics are divided into those that assess cell type alignment performance and those that measure effectiveness of batch effect correction [10]. Overall score takes the weighted average over cell type matching score and batch correction score $(0.6 * \text{Avg. Bio}) + (0.4 * \text{Avg. Batch})$. For certain models results have a small amount of variance, either because the model is fine tuned model (scVI, scArches), or because the model includes sampling (UCE). For these models, the dataset is shuffled using 10 different random seeds, and then the method is scored. For the fine tuned models, a new model was trained on the shuffled data for each seed.

| Query Cell Type (Tabula Sapiens) | Nearest Neighbor 1 | Nearest Neighbor 2 | Nearest Neighbor 3 |
|---|---|---|---|
| **tuft cell of colon** | brush cell | intestinal tuft cell | tuft cell of colon |
| **intestinal crypt stem cell of colon** | transit amplifying cell of colon (TS) | paneth cell of colon (TS) | colon epithelial cell |
| **monocyte** | myeloid dendritic cell (TS) | myeloid cell | colon macrophage (TS) |
| **myofibroblast cell** | smooth muscle cell | kidney interstitial cell | tracheobronchial smooth muscle cell |
| **colon macrophage** | myeloid dendritic cell (TS) | myeloid leukocyte (TS) | myeloid cell |
| **natural killer cell** | mature nk t cell (TS) | gamma-delta t cell (TS) | cd8-positive, alpha-beta t cell (TS) |
| **plasma cell** | IgA plasma cell | plasma cell | IgG plasma cell |
| **myeloid dendritic cell** | colon macrophage (TS) | myeloid leukocyte (TS) | myeloid cell |
| **enterocyte of epithelium of large intestine** | colon epithelial cell | best4+ intestinal epithelial cell, human (TS) | large intestine goblet cell (TS) |
| **transit amplifying cell of colon** | intestinal crypt stem cell of colon (TS) | paneth cell of colon (TS) | transit amplifying cell |
| **myeloid leukocyte** | colon macrophage (TS) | myeloid dendritic cell (TS) | myeloid cell |
| **fibroblast** | fibroblast | stromal cell | myofibroblast cell |
| **mast cell** | mast cell | granulocyte | basophil |
| **innate lymphoid cell** | mature nk t cell (TS) | cd4-positive, alpha-beta t cell (TS) | natural killer cell (TS) |
| **paneth cell of colon** | intestinal crypt stem cell of colon (TS) | transit amplifying cell of colon (TS) | best4+ intestinal epithelial cell, human (TS) |

**Supplementary Table 6: Tabula Sapiens v2 Cell Type Alignments to IMA for Large Intestine Tissue.** The first column corresponds to the query cell type from Tabula Sapiens v2 that is mapped zero-shot to the IMA UCE embedding. The 3 columns after that are nearest neighbor cell type centroids to the query in the UCE embedding space. The '(TS)' notation represents a cell type that was present in the mapped data from Tabula Sapiens v2. We treat this as an incorrect match since it would indicate the presence of a dataset-specific effect where a different cell type from the same experiment was mapped closer to the query cell type than a correctly matching cell type in the IMA (Methods). Green indicates a correct match, Red indicates an incorrect match and Yellow indicates a match that is correct at coarser resolution.

| Query Cell Type (Tabula Sapiens) | Nearest Neighbor 1 | Nearest Neighbor 2 | Nearest Neighbor 3 |
|---|---|---|---|
| **smooth muscle cell** | smooth muscle cell | kidney interstitial cell | tracheobronchial smooth muscle cell |
| **macrophage** | myeloid cell | monocyte | macrophage |
| **neutrophil** | neutrophil | bone marrow cell | myeloid cell |
| **luminal cell of prostate epithelium** | luminal cell of prostate epithelium | epithelial cell of prostate | basal cell of prostate epithelium (TS) |
| **endothelial cell** | vein endothelial cell | blood vessel endothelial cell | endothelial cell |
| **erythrocyte** | enucleated reticulocyte | enucleate erythrocyte | reticulocyte |
| **basal cell of prostate epithelium** | conjunctival epithelial cell | luminal cell of prostate epithelium (TS) | epithelial cell of prostate |
| **fibroblast** | alveolar type 2 fibroblast cell | fibroblast | kidney interstitial fibroblast |
| **mast cell** | mast cell | basophil | granulocyte |
| **b cell** | memory B cell | naive B cell | B cell |
| **mature nk t cell** | natural killer cell | mature NK T cell | CD16-positive, CD56-dim natural killer cell, human |
| **t cell** | erythrocyte (TS) | enucleated reticulocyte | erythrocyte |
| **cd4-positive, alpha-beta t cell** | cd8-positive, alpha-beta t cell (TS) | T cell | effector CD4-positive, alpha-beta T cell |
| **cd8-positive, alpha-beta t cell** | cd4-positive, alpha-beta t cell (TS) | activated CD8-positive, alpha-beta T cell | T cell |

**Supplementary Table 7: Tabula Sapiens v2 Cell Type Alignments to IMA for Prostate Tissue.** The first column corresponds to the query cell type from Tabula Sapiens v2 that is mapped zero-shot to the IMA UCE embedding. The 3 columns after that are nearest neighbor cell type centroids to the query in the UCE embedding space. The '(TS)' notation represents a cell type that was present in the mapped data from Tabula Sapiens v2. We treat this as an incorrect match since it would indicate the presence of a dataset-specific effect where a different cell type from the same experiment was mapped closer to the query cell type than a correctly matching cell type in the IMA (Methods). Green indicates a correct match, Red indicates an incorrect match and Yellow indicates a match that is correct at coarser resolution.

| Original Species | What species's protein embeddings are used? | Loss ↓ | SCIB Bio Conservation Score ↑ |
|---|---|---|---|
| Mouse | Mouse | **0.398** | **0.629** |
| Mouse | Human | 0.490 | 0.538 |
| Human | Human | **0.421** | **0.549** |
| Human | Mouse | 0.522 | 0.506 |

**Supplementary Table 8: Homolog Gene Performance for Primary Motor Cortex.** Model loss and Bio Conservation score are evaluated on human and mouse primary cortex[34]. First, both human and mouse datasets are subset to a common set of one to one homologous genes. Then, UCE's performance on each species is evaluated, using either the species' original genes, or the other species' genes. Switching between genes is done by using the other species protein embeddings. For both human and mouse, using homologous genes from the other species degraded performance.

| Original Species | What species's protein embeddings are used? | Loss ↓ | SCIB Bio Conservation Score ↑ |
|---|---|---|---|
| Mouse | Mouse | **0.408** | **0.506** |
| Mouse | Human | 0.454 | 0.498 |
| Human | Human | **0.421** | **0.573** |
| Human | Mouse | 0.481 | 0.558 |

**Supplementary Table 9: Homolog Gene Performance for Embryonic Limb Atlas.** Model loss and Bio Conservation score are evaluated on the human and embryonic limb atlas[35]. First, both human and mouse datasets are subset to a common set of one to one homologous genes. Then, UCE's performance on each species is evaluated, using either the species' original genes, or the other species' genes. Switching between genes is done by using the other species protein embeddings. For both human and mouse, using homologous genes from the other species degraded performance.

| Embedding Method | Isolated labels | KMeans NMI | KMeans ARI | Silhouette label | cLISI | Bio conservation |
|---|---|---|---|---|---|---|
| Geneformer | 0.464 | 0.282 | 0.039 | 0.394 | 0.990 | 0.434 |
| scGPT | 0.569 | 0.596 | 0.141 | 0.483 | 0.997 | 0.557 |
| UCE | **0.591** | 0.703 | 0.255 | 0.508 | **1.000** | 0.611 |
| PCA | 0.589 | 0.644 | 0.200 | 0.454 | 0.998 | 0.577 |
| scVI | **0.591** | **0.710** | **0.263** | **0.509** | 0.998 | **0.614** |

**Supplementary Table 10: Performance on Human Brain Cell Atlas** Model performance on the Human Brain Cell Atlas[9] is evaluated against other methods in the zero-shot setting, and to fine tuned methods. Bio Conservations scores are calculated for the cluster_id cell type label, containing 382 cell types, and are reported as the mean score for 10 independent subsamples of 500,000 cells each (Supplementary Note 7). PCA and scVI are computed on 5,000 highly variable genes, calculated from the whole dataset. Bolded numbers represent the highest score for a given metric, while underlined numbers represent the second highest score.

| Comparison Condition | Mean Cosine Similarity | Empirical Standard Deviation | Interpretation |
|---|---|---|---|
| **Cell vs. Itself** (Different Random Seed) | 0.957 | 0.032 | High Stability: the embedding is robust to sampling noise |
| **Cell vs. Same Cell Type** | 0.741 | 0.088 | Biological Cluster: distinct from self, biologically grouped |
| **Cell vs. Random Cell** | 0.335 | 0.165 | Background: expected distance between unrelated cells from same dataset |

**Supplementary Table 11: Effect of Gene Sampling on Cell Embedding Similarity** To assess the impact of the model's random sampling of genes on embedding stability, similarity of cells' embeddings are measured for a subsample of 100,000 cells from Tabula Sapiens [30]. The embedding of a cell is most close to itself in the embedding space when a different random seed is used (Cell vs. Itself), very close to other cells from the same cell type (Cell vs. Same Cell Type), and the least similar to other random cells.

| **Dataset: Tabula Sapiens v1 and v2** | | | |
|---|---|---|---|
| **Method** | **Rank-PCA** | **Corr-PCA** | **Corr-Weighted** |
| **UCE** | **0.221** | **0.534** | **0.331** |
| **scGPT** | 0.197 | 0.305 | 0.166 |
| **Geneformer** | 0.138 | 0.239 | 0.063 |
| **Dataset: Human Brain Cell Atlas** | | | |
| **Method** | **Rank-PCA** | **Corr-PCA** | **Corr-Weighted** |
| **UCE** | **0.564** | **0.652** | **0.673** |
| **scGPT** | 0.475 | 0.528 | 0.570 |
| **Geneformer** | 0.210 | 0.211 | 0.247 |

**Supplementary Table 12: Performance on scGraph Benchmark** We assess the performance of UCE, scGPT and Geneformer using the scGraph [36] benchmark (version 0.1.2). scGraph measures how well cell embedding spaces consistently capture cell type relationships across different samples. We perform the benchmark on Tabula Sapiens v1 and v2 (1,194,951 cells and 180 cell types, 534 batches) [30], and the Human Brain Cell Atlas (2,480,956 cells, 382 cell types, 606 batches) [9]. UCE outperforms scGPT and Geneformer on all metrics on both datasets. To calculate the PCA values for each dataset, expression counts are normalized to the median value and then log transformed using defaults, followed by standard scanpy PCA calculation with 50 principal components.

| Gene | Log2 Fold Change IPF vs COPD | Adjusted P Value IPF vs COPD | Log2 Fold Change IPF vs Control | Adjusted P Value IPF vs Control |
|---|---|---|---|---|
| *Dcn* | -0.123 | 0.713 | -0.352 | 0.116 |
| *Lpar1* | 0.045 | 0.915 | -.160 | 0.588 |
| *Col1a1* | 1.515 | 0.003 | 2.000 | 0.000005 |
| *Cxcl14* | 0.910 | 0.045 | 1.481 | 0.00005 |
| *Cxcl12* | 0.136 | 0.800 | 1.367 | 0.000003 |
| *Col5a1* | 1.093 | 0.0008 | 1.148 | 0.00003 |
| *Col5a2* | 0.775 | 0.003 | 1.173 | 0.00000008 |
| *Col1a2* | 1.156 | 0.002 | 1.317 | 0.00003 |
| *Col3a1* | 1.638 | 0.0008 | 1.521 | 0.0003 |
| *Egln1* | -0.790 | 0.032 | -0.641 | 0.0533 |

**Supplementary Table 13: Norn Cells Pseudobulk Differential Expression** Predicted norn-like cells in patients with COPD, control and IPF are aggregated at the patient identity level using decoupler version 2.1.4 and filtered with min_cells= 10 and min_counts= 1000. Next, differential expression is calculated using DefaultInference (wald-test) implemented in pydesec2 version 0.5.4 for IPF patients' predicted norn cells versus COPD and control patients'.

| Dataset | UCE | SATURN | SAMap |
|---|---|---|---|
| **Green Monkey Lung** | 0.766 | **0.945** | 0.681 |
| **Naked Mole Rat Spleen** | **0.652** | 0.122 | 0.640 |
| **Chicken Retina** | **0.761** | 0.280 | 0.565 |
| **Chicken Heart** | 0.330 | 0.300 | **0.464** |

**Supplementary Table 14: Novel Species Label Transfer Performance** We assess the performance of UCE, SATURN and SAMap on the task of transferring labels from a tissue-matched reference human dataset to a new species (Supplementary Note 5). The scores report the accuracy of the predicted labels for the novel species. Importantly, SATURN and SAMap are fine tuned methods, and yet, UCE still outperforms each method on 3 out of 4 of the datasets, even though it is in the more challenging, zero-shot setting.
